# Distinct evolutionary patterns of tumor immune escape and elimination determined by ECM architectures

**DOI:** 10.1101/2024.05.13.594017

**Authors:** Yijia Fan, Jason T. George

## Abstract

Cancer progression remains a significant clinical challenge. Phenotypic adaptation by tumor cells results in disease heterogeneity, which drives treatment resistance and immune escape. T cell immunotherapy, while effective at treating some cancer subtypes, can also fail due to limits on tumor immunogenicity or T cell recognition. For example, one potential contributor to immune escape involves the density and alignment of the extracellular matrix (ECM) surrounding tumors, also known as Tumor-Associated Collagen Signature (TACS). However, the specific mechanisms by which aligned fibers contribute to decreased patient survival rates have not yet been decoupled. Here, we developed our EVO-ACT (EVOlutionary Agent-based Cancer T cell interaction) model to study how TACS affects tumor evolution and dynamic tumor-T cell interactions. We identified a variety of TACS-specific dynamical features that influence T cell infiltration, cancer immunoediting, and ultimate immune escape. Our model demonstrates how TACS and phenotypic adaptation together explain overall survival trends in breast cancer.

## Introduction

The immune system plays a central role in the adaptive response against tumor progression, wherein cytotoxic (CD8+) T cells attempt to engage with and eliminate the cancer population. This interaction results in immunoediting of tumor populations that can either lead to tumor escape, elimination, or a sustained equilibrium period (1–5). Prior experimental and theoretical work has been directed at understanding how repeated tumor-immune interactions affect the ultimate dynamics of cancer progression and escape (6–10). These earlier models have described how clonally heterogeneous cancer populations evolve under adaptive immune selective pressures.

It is now appreciated that the adaptive immune system can capably clear cancer in some cases, while in others tumor immune escape occurs. The immune microenvironment plays an important and multifaceted role in this process. One unanswered question relates to the role of ECM organization and its effects on tumor immune recognition. In solid malignancies, ECM geometry in the microenvironment has been associated with disease stage (11, 12) and observed T cell infiltration (13, 14). Specifically, empirically observed ECM topologies are frequently categorized based on fiber arrangement: random fibers (TACS1), circumferentially aligned fibers (TACS2), and radially arranged fibers (TACS3), illustrated in Figure 1 (12). Despite the fact that a clear negative correlation between TACS and patient survival has been established (15), the specific roles and extent of TACS in sculpting T cell-driven cancer evolution remain uncharacterized. Mechanisms underlying how TACS influences cell movement are still not fully elucidated, with divergent opinions on the precise details involving how and to what extent the ECM mediates immune cell infiltration (13, 14, 16– 18). At present, we currently lack a physical model relating the impact of TACS on the spatial co-evolution between an adaptive immune repertoire and a heterogeneous population of evading cancer cells.

**Fig. 1.**
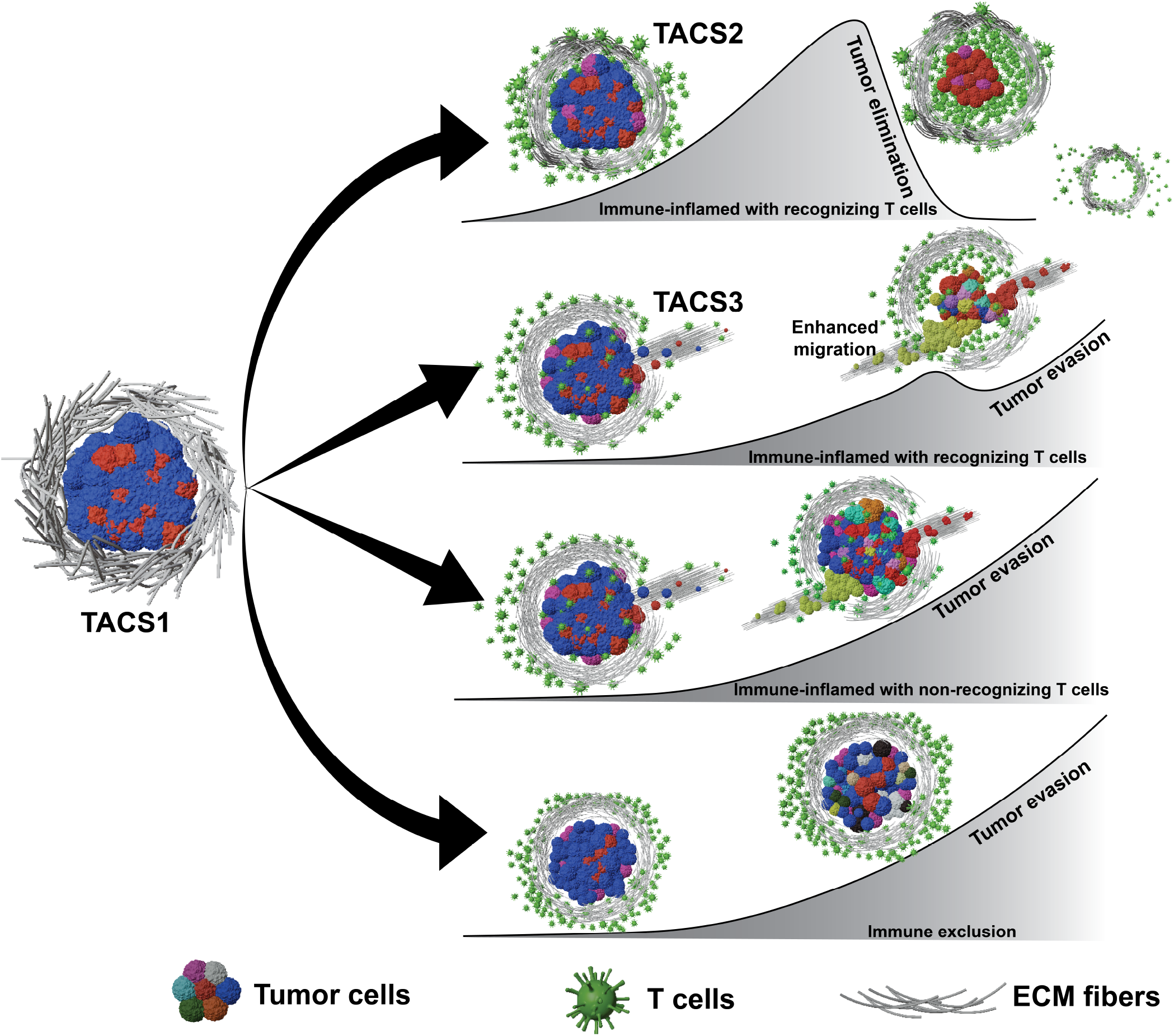
Three common TACS architectures and immune infiltration scenarios. EVO-ACT considers tumors initially in TACS1, which then progress to TACS2 and TACS3. Interactions between T cells and cancer cells can give rise to tumor elimination, temporary equilibrium, or escape. Tumor heterogeneity arises through stochastic adaptation resulting in variable TAA presentations.

To address this, we develop and apply our EVO-ACT model to study how TACS influences tumor evolution and the dynamic interaction between tumor and T cells. We quantify differences in the dynamics of cell migration and evasion, in addition to the spatial distributions of cancer cells and immune cells, as a functional consequence of TACS. Our results suggest that the degree of cancer immunoediting is dependent on TACS-specific differences in T cell infiltration and moving efficiency, and that TACS have a greater impact on chemokine-directed T cell infiltration than they do on evading tumor cell evasion. When applied to predict differences in TACS3-dependent disease progression, we find that our modeling framework requires the inclusion of additional phenotypic adaptation mechanisms, such as the Epithelial-to-Mesenchymal Transition (EMT), in order to successfully recapitulate clinically observed cancer survival trends. Our model predicts that immunogenicity differences via decreased Tumor-Associated Antigen (TAA) availability and immune checkpoint upregulation synergize to result in immune escape, which successfully predicts overall survival trends in breast cancer (19, 20). The EVO-ACT framework provides a detailed dynamical description of the role of TACS in tumor evolution when subject to adaptive immune selective pressure. We anticipate that its use can be more broadly applied to understand cancer evolutionary patterns and treatment success or failure in specific cases where observed TACS architecture and phenotypic status are previously defined.

## Results

### Stochastic agent-based modeling

#### Cancer cells

To model cancer cells, we assumed that a small collection (*n ∼* 4000) corresponding to a total tumor diameter of 0.1cm of tumor precursors reside at the center of a circular Region of Interest (ROI), from which active tumor cells may extravasate outward, invade, and divide. These tumor cells, together with CD8+ T cells, comprise agents in our model. We assumed that cancer cells may divide and migrate at percell rates of *λ* and *α*_*t*_, respectively. Cancer cells are also assumed to possess an initial collection of shared TAAs displayed on their surface. For foundational understanding, we generated TAAs randomly according to a Poisson (*ν* = 100) distribution. Each cell may also undergo mutation or phenotypic adaptation with a rate of *μ* = 5 *·*10^−4^ per cell division. We assumed that each adaptation event results in an equal likelihood of the addition or removal of a Poisson number of TAAs with a mean of 10, and that each TAA addition or removal increases or decreases the division rate by 1% of *λ* with equal probability. Evolution in this model results in distinct cancer populations, or “clones”, which are distinguished only based on their expression of TAAs and may occur through genetic or epigenetic adaptation mechanisms. We accounted for the “contact guidance” theory of migration by assuming that each cancer cell randomly selects a fiber within 5 cell diameters (75*μ*m) along which to migrate (12), and equal probabilities are given to both fiber directions. We also considered variability in the starting division rates and TAA abundances.

In simulations that incorporate EMT, we assume that EMT occurs once a certain total tumor burden is achieved. Upon reaching this threshold, all dividing cells at the tumor periphery are assumed to undergo EMT, which was modeled by a decreased division rate (five-fold reduction), increased migration rate (five-fold increase), and decreased immunogenicity (reduction of TAAs by a Poisson-distributed random variable with mean of 15) to match previous experimental observations (20, 21).

We utilized a Gillespie simulation-based approach to model cancer division and migration dynamics.

#### CD8+ T cells

T cells comprise the other active agents in our modeling framework, and we modeled their dynamics by starting with an initial population of (*N* = 5000) T cells located at the boundary of the ROI. These T cells migrate inward at a deterministic rate of *α*_*T*_. Their directed migration is influenced by the surrounding extrinsic microenvironment, which includes collagen fiber alignment and chemotaxis (12, 22). To model differential antigen specificities of distinct T cell clones, we assumed that distinct T Cell Receptors (TCR) exist (*R*_*n*_ *≤*5000), each having the capability to recognize various antigen signatures. In this model, T cell diversity is characterized by the absolute number of distinct antigen specificities the T cell population can cover, which correlates directly with the number of TAA-specific clones (*r*_*n*_). T cell recognition is thus a function of both *R*_*n*_ and *r*_*n*_. To model recognition, each migrating T cell surveys the vicinity (*ϵ*_*k*_ = 45 *μ*m). Upon encountering at least one recognizable TAA, T cells eliminate the tumor cell and subsequently divide. To account for lineage-specific T cell contractions in the absence of antigen signaling, daughter T cell possess a finite survival window, after which they are removed if they cannot recognize and eliminate another cancer cell (see Figure S2 for full details). While we do not explicitly include memory T cells in our model, our assumptions on T cell dynamics maintain a small population of effector T cell clones that have previously recognized TAAs.

#### ECM topology

We modeled ECM fibers situated outside the tumor core with normally distributed lengths (mean *μ* = 10*μ*m; variance *σ*^2^ = *μ/*10) (23). Fiber density was assumed to decrease radially outward from the tumor center, and initially all fibers are randomly oriented, corresponding to TACS1 (24). To account for tumor-driven TACS remodeling (12), we allowed for each dividing tumor cell at the periphery to alter the fiber direction from TACS1 to TACS2 within a neighborhood of the cell (5 cell diameter, 75*μ*m). Based on previous findings, tumor cells can remodel fibers into a radial pattern within a 5-fold neighborhood relative to tumor radius (25). For simplicity, based on the size of the central tumor, we assumed that dividing cells at the tumor boundary will collectively remodel fibers within *∼*0.18 cm into TACS3. We also studied the role of heterogeneous fiber alignment by adjusting the variance of fiber orientation in each of TACS2 and TACS3. We refer the reader to Figure S1 for the initial spatial distribution of tumor, T cells, and fibers in our model and the Methods section for full model details.

### Cancer evolution generates TACS-specific spatial signatures in the absence of immune selective pressure

In our model, tumor populations grow and acquire alterations that enhance their division rate, but also affect their antigenic burden. In the absence of T cell recognition, cancer cells exhibited evolutionary trajectories comprised of outward growth, and clones with increased fitness through enhancements in their growth rates are positively selected. Figure 2A represents one stochastic realization of this process for a single tumor population that divides until reaching a diameter of *∼* 0.5 cm. Our results capture the spatial variability in tumor heterogeneity and are in qualitative agreement with prior models in the regime of tumor growth and invasion into neighboring tissue (26).

**Fig. 2.**
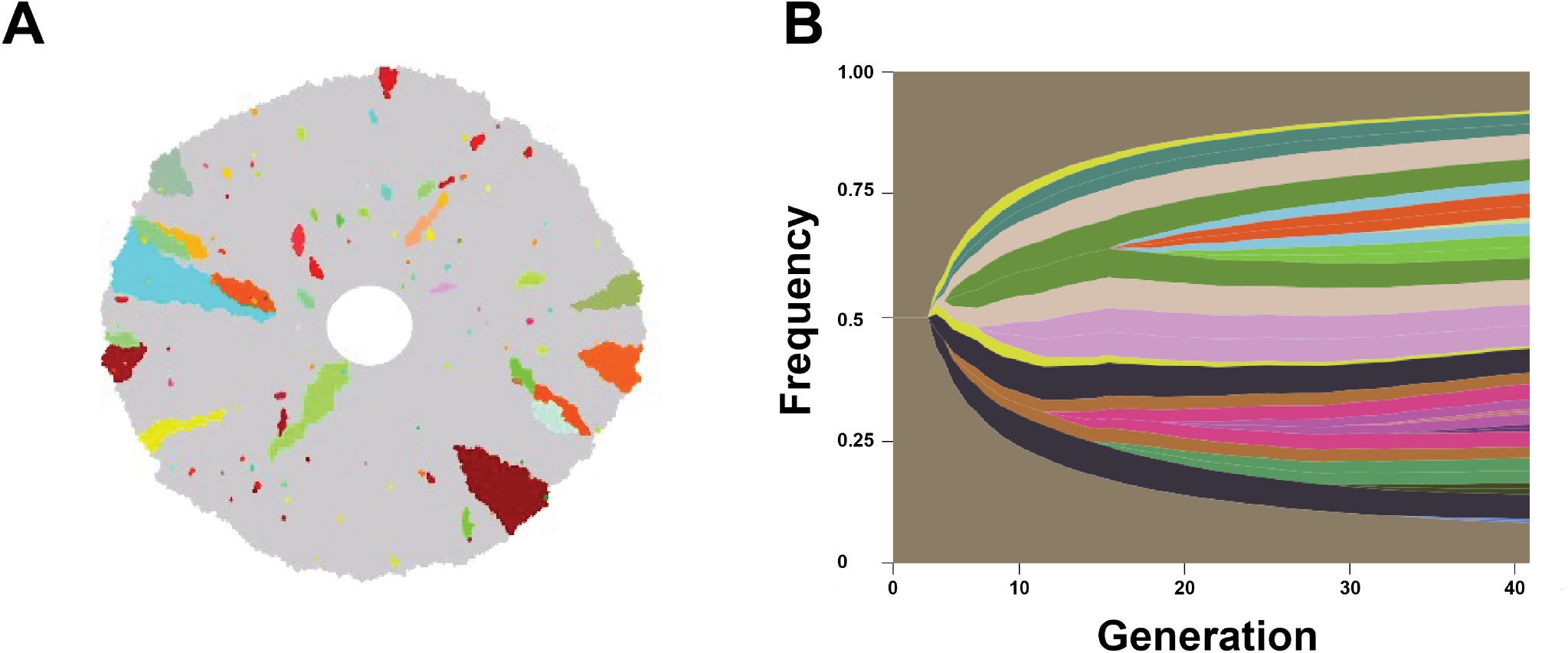
Spatial heterogeneity obtained through cancer evolutionary progression. A: The spatial distribution of cancer clones is illustrated following progression to ∼ 0.5cm diameter (corresponding to *∼*110000 tumor cells). Gray cells represent daughter cells arising from the founder clone, and additional colors distinguish subsequent clones that harbor distinct antigen expression patterns. B: A representative fish plot indicating the frequency of each antigenic subclone, where the brown represents the initial clone.The vertical axis represents the frequency of each tumor clone and the horizontal axis represents the generation.

Given the well-established clinical relevance of TACS (11, 12, 15, 27–29), we next aimed to identify how TACS impacts cancer patient survival or tumor elimination and escape. To investigate this, we first explored how TACS influences the encounter between tumors and T cells. Based on the “contact guidance” theory (30), we hypothesized that TACS affects the spatial distribution and migration of cells. To test this, we simulated an initially small collection of tumor cells dividing within the three types of TACS in the absence of immune pressure, with TACS2 and TACS3 being perfectly aligned. For each TACS architecture, a distinct pattern of tumor spatial distribution emerged: tumor cells are randomly packed in TACS1, they are tightly encircled in TACS2, and radially arranged in TACS3 (Figure 3A). These patterns also manifest in corresponding random (TACS1), circumfrential (TACS2), and radial (TACS3) spatial distributions of tumor clones (Figure 3A). Subsequent analysis of single-cell cancer migration trajectories also revealed differences in their motion, with a random-walk pattern in TACS1, an outward “zig-zag” motion in TACS2, and an “outward radiating” motion in TACS3. Together, these results highlight how distinct TACS can significantly influence cancer cell migration and the subsequent spatial heterogeneity observed across otherwise identical underlying tumor evolutionary processes. We postulated that such differences may be relevant for immune recognition since, for example, the surface area of the tumor boundary in TACS3 is substantially larger than TACS2 with fewer subclones protected in the tumor interior.

**Fig. 3.**
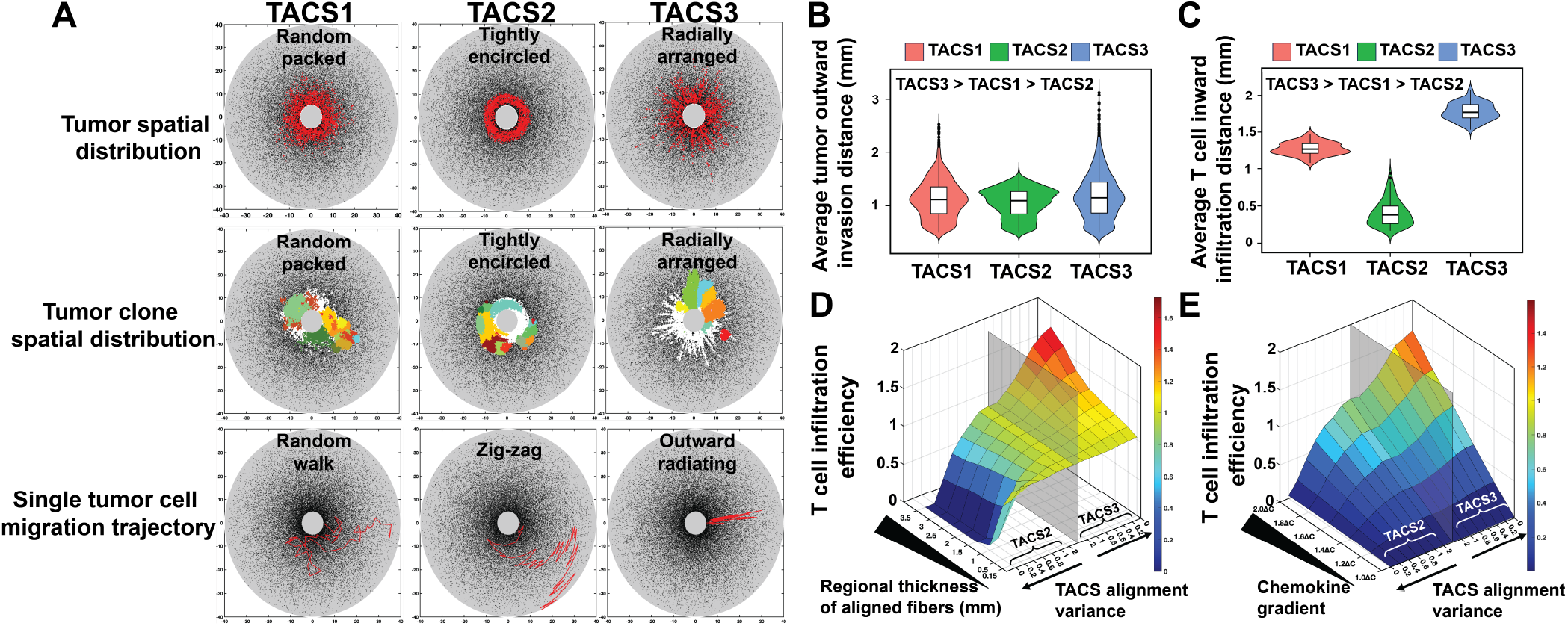
TACS influences spatial distribution, migration direction and efficiency of both tumor cells and T cells, with chemical attractant amplifying TACS impact on T cell infiltration efficiency. A: Tumor spatial distribution (λ= 0.1, α_*t*_ = 0, α_*T*_ = 0), antigenic clonal variant spatial distribution (λ= 0.1, α_*t*_ = 0, α_*T*_ = 0) and single-cell migration trajectories (λ= 0, α_*t*_ = 0.1, α_*T*_ = 0) in TACS1-3. B: Average tumor invasion distance (*n* = 10^3^ tumor cells) across TACS1 to TACS3 within a fixed time frame. TACS2 and TACS3 are perfectly aligned. λ= 0, α_*t*_ = 0.1, α_*T*_ = 0. C: Average T cell infiltration distance (*N* = 10^3^ T cells) across TACS1 to TACS3 within the consistent time frame depicted in B. TACS2 and TACS3 are perfectly aligned. λ= 0, α_*t*_ = 0, α_*T*_ = 0.1 D: Normalized T cell infiltration efficiency in TACS1-3 with varying alignment region thickness and alignment variance. We measured the time taken for N=500 T cells to infiltrate from the boundary of the ROI to the boundary of the central tumor circle under varying thicknesses of the alignment region and varying alignment variances of TACS2 and TACS3. This time was normalized against the time spent in TACS1. The semi-transparent gray plane in the graph represents the baseline or T cell infiltration efficiency in TACS 1. The left side of this plane represents the infiltration efficiency of T cells in TACS2, with TACS2 features becoming more pronounced as one moves further to the left. Conversely, the right side of this plane represents the infiltration efficiency of T cells in TACS3, with TACS3 features becoming more pronounced as one moves further to the right. The y-axis represents the thickness of the alignment region, with thicker regions positioned toward the inner side. λ= 0, α_*t*_ = 0, α_*T*_ = 0.1. E: Normalized T cell infiltration efficiency in TACS1-3 with varying chemokine gradients and alignment variances. We replicated the experiments in D, with a default aligned region thickness of 0.5mm, and replaced the regional thickness of aligned fiber with chemokine gradients. λ = 0, α_*t*_ = 0, α_*T*_ = 0.1.

### TACS generates distinct patterns of cell migration and consequent cancer immunoediting

#### TACS determines cancer and T cell migration efficiencies

To quantify the role of TACS on cancer and T cell migration, we next tracked migration by simulating (*n* = 10^3^) tumor cells undergoing random migration and (*N* = 10^3^) T cells undergoing directed migration toward a chemokine signal over a fixed time period. In each case, we maintained perfect alignment of TACS2 and TACS3 for comparison purposes. Our simulations predict that migration efficiencies in our model are maximal in TACS3, intermediate in TACS1, and minimal in TACS2 for both T cells and tumor cells (Figure 3B-C). These findings are consistent with the fact that radially oriented TACS3 fibers direct T cell movement to the primary tumor mass, while TACS2 results in circumferential movement that reduces the overall migration rate observed in randomly oriented TACS1 fibers. Despite similar trends in the relative cancer and immune cell movements for each TACS, these differences were substantially more pronounced for T cells than for cancer cells (Figure 3B-C). These results suggest that the benefit to migration of TACS3 significantly favors T cell infiltration over cancer escape.

To better understand the role of TACS fibers and cell signaling on T cell migration, we performed additional simulations tracking the migration efficiency of (*N* = 500) T cells from the ROI boundary to the tumor center boundary under TACS1-3. We also independently varied the regional thickness of aligned fibers and the strength of the chemokine gradient guiding T cell migration. Migration efficiency was taken to be the inverse of mean migration time, and each result is normalized to the mean values obtained for TACS1. Our results (Figure 3D-E) illustrate the spectrum on which highly vs. loosely aligned TACS influence T cell migration. Larger regions of TACS fibers amplify the observed TACS-specific migration differences, and a larger chemokine gradient consistently enhances infiltration efficiency across all TACS conditions. Moreover, our results show that TACS2 does not constitute an absolute barrier to T cell infiltration; rather, it diminishes the efficiency of T cell infiltration. The degree to which this efficiency is diminished relies on factors such as the degree of alignment and thickness of the TACS2 region. Our findings support previous work showing that clinically observed patterns of T cell exclusion are consistent with differences in T cell chemical signaling rather than a physical barrier to T cell infiltration (17).

#### TACS-specific tumor spatial heterogeneity and T cell infiltration together generate variety in cancer immunoediting

We next sought to understand how TACS-dependent spatial distributions in cancer clones and variable T cell infiltration efficiencies together affect tumor elimination and escape. Given that T cell infiltration is often correlated with greater patient overall survival across various tumor subtypes (31–33), we first quantified the extent of T cell infiltration in TACS environments by comparing the spatial distribution of T cell density in the tumor core and tumor margin. Since our model does not include other cell types present in the tumor microenvironment (TME), we assumed that the tumor margin in our model is immediately exterior to the boundary of the tumor core and extends outward by 250 *μ*m (17). We performed largescale stochastic simulations of tumor growth, T cell infiltration and eventual tumor-T cell interactions for TACS3^−^ (Figure 4A-D) and TACS3^+^ (Figure 4E-H) conditions. In each case, we calculated an average T cell density in the tumor core relative to the tumor margin to differentiate infiltrating and non-infiltrating T cells at the tumor boundary (indicated by the red dashed lines in Figure 4B,D,F,H).

**Fig. 4.**
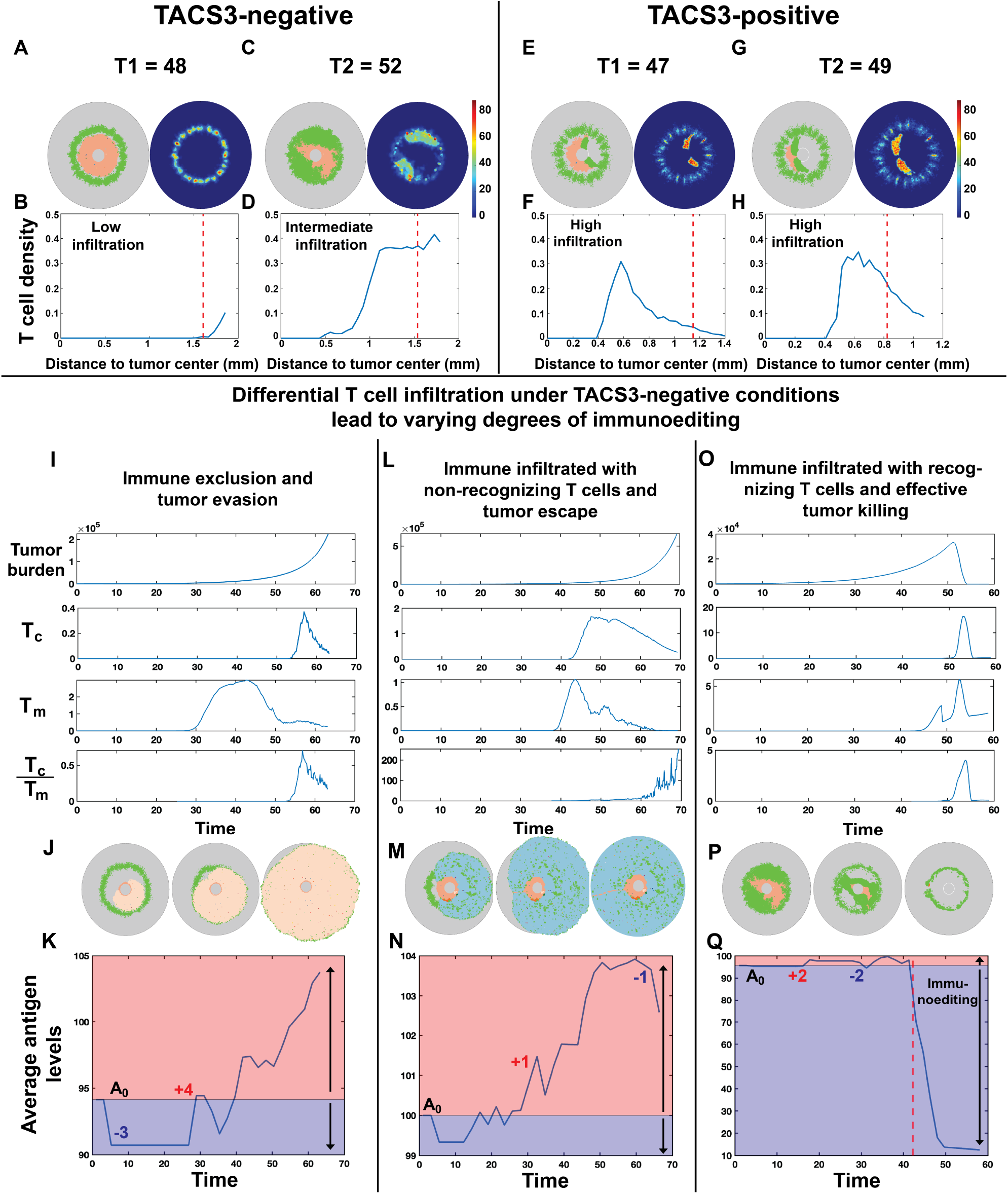
TACS impacts spatial heterogeneity and infiltration levels of T cells, consequently influencing immunoediting. TACS3 promotes more efficient T cell infiltration. A, C, E, G: Illustrative examples of tumor-T cell interaction snapshots and associated T cell density heatmaps within TACS3^−^ (A, C) and TACS3^+^ (E, G) conditions. B, D, F, H: The radial distribution of T cell density from the tumor center to the boundary of ROI is depicted. The red dashed line represents the boundary of the tumor core in the above two conditions, *R*_*n*_ = 210, *r*_*n*_ = 10. I, L, O: Tumor burden and T cell density were quantified in the tumor core (*T*_*c*_) and margin (*T*_*m*_), and *T*_*c*_:*T*_*m*_. In immune exclusion and non-recognizing T cell inflammation conditions, *R*_*n*_ = 210, *r*_*n*_ = 0. In recognizing T cell inflammation condition, *R*_*n*_ = 210, *r*_*n*_ = 10. J, M, P: Snapshots of tumor and T cell interactions in three T cell infiltration conditions. K, N, Q: The average antigen levels were analyzed for each of the three conditions across 10 iterations. In each plot, initial antigen levels (*A*_0_) partition the graph into red (antigen gain) and blue (antigen loss) segments. Higher y-values indicate higher antigenicity. The red dashed line in Q marks the onset of widespread tumor killing or immunoediting by T cells. In all conditions, λ =0.1, α_*t*_=0.02, α_*T*_ =1.8.

We observed distinct profiles of T cell infiltration in TACS3^−^ (Figure 4A-D) and TACS3^+^ (Figure 4E-H) conditions. Heat maps depicting T cell densities indicate regions where active T cell recognition and concomitant expansion occurs (Figure 4A,C,E,G). In particular, TACS3^+^ environments are characterized by high T cell infiltration into the tumor core relative to the tumor margin. These findings contrast with TACS3^−^ environments, which were characterized by low to intermediate levels of T cell infiltration (Figure 4B,D,F,H). These trends were maintained in additional simulations that assumed perfectly aligned TACS2 and TACS3 signatures (Figure S3). The observation that enhanced infiltration and consequent T cell recognition occurs in a TACS3 environment is consistent with the prior finding that TACS3 provides greater benefit to T cell mobility (34, 35). Moreover, reductions in T cell infiltration in TACS2 still permit T cell recognition and do not function as an absolute barrier to recognition, even in perfectly circumscribed tumors (Figure S3). Lastly, greater TACS2 thickness enhances the inhibitory capacity on T cells, thereby promoting tumor evasion (Figure S4).

Given these differences in T cell infiltration, we next sought to observe how TACS-specific effects on cell migration and tumor heterogeneity together impact immunogenicity during tumor progression. We performed additional simulations to assess TACS3^−^ T cell infiltration in greater detail. We first modeled an immune-excluded phenotype by confining 95% of T cells to the tumor margin. These simulations were compared against cases that allowed for T cell infiltration. In all conditions, we tracked T cell infiltration into the tumor core (*T*_*c*_), tumor margin (*T*_*m*_), and the ratio *T*_*c*_ : *T*_*m*_ (Figure 4I,L,O). We subsequently repeated 10 replicates for each condition, and average tumor antigenic burden was calculated over time in each case (Figure 4K,M,Q). TACS3^−^ cases resulted in both immune-excluded and immune-infiltrated tumors (Figure 4I-Q), and only a subset of cases with T cell infiltration exhibited significant T cell expansions, which we distinguish as ‘non-recognizing’ (Figure 4L-N) or ‘recognizing’ (Figure 4O-Q). In both immune-excluded and infiltrated non-recognizing cases, we observed a steadily increasing average antigenic burden during cancer progression, which is comparable to empirically observed mutation accumulation rates (Figure 4K,N) (36, 37). This behavior contrasted with the infiltrated recognizing case, wherein TAA availability notably declined over time and is consistent with strong immunoediting (Figure 4O-Q). Our model predicts that active immune pressure results in tumor clones that tend to become less immunogenic over time, suggestive of the strong selective pressure operant during T cell infiltration and in agreement with previous findings (2, 38). Moreover, these dynamics underscore the fact that T cell infiltration alone is insufficient in discerning T cell response, with spatially dependent T cell expansion more indicative of recognition and tumor elimination. Figure S5 illustrates how TACS indirectly leads to different expansion patterns of T cells by affecting the spatial positioning of tumor clones or TAAs.

### TACS3-associated phenotypic adaptability decreases predicted patient survival rates and impairs the efficiency of checkpoint inhibitor therapy

#### TACS3-associated phenotypic changes significantly diminish survival rates

Our previous results suggested that TACS3 in isolation confers an overall net benefit to immune recognition (Figure 3C). This prediction was unexpected in light of the fact that survival rates in TACS3^+^ conditions are generally lower than those observed for TACS3^−^ conditions (15).

To investigate this further, we performed largescale stochastic simulations in perfectly aligned TACS2 and TACS3 conditions, each characterized by varying TCR recognition diversity. Tumor death was assumed at ∼0.5 cm (*∼*110,000 cells), with recorded times. We explored several parameter regimes but consistently found that TACS3 conditions resulted in enhanced survival compared to TACS2 cases (Figure 5A). These findings suggest that TACS-specific changes alone do not explain observed survival in our modeling framework, and they motivated us to account for additional phenotypic changes that are known to be present in later diseases. To study this, we introduced EMT, which is a commonly occurring transition in epithelial cancers during disease progression (39). By introducing EMT separately in TACS2 and TACS3, we observed higher survival rates in TACS3 compared to TACS2 (Figure 5B, Figure S6A). Our simulations that include EMT only after TACS2-TACS3 progression most closely match clinically observed (15) outcomes. (Figure 5C). These findings are further supported by experimental evidence demonstrating that ECM influences EMT regulation, and reciprocally, EMT induces changes in ECM remodeling (40–42).

**Fig. 5.**
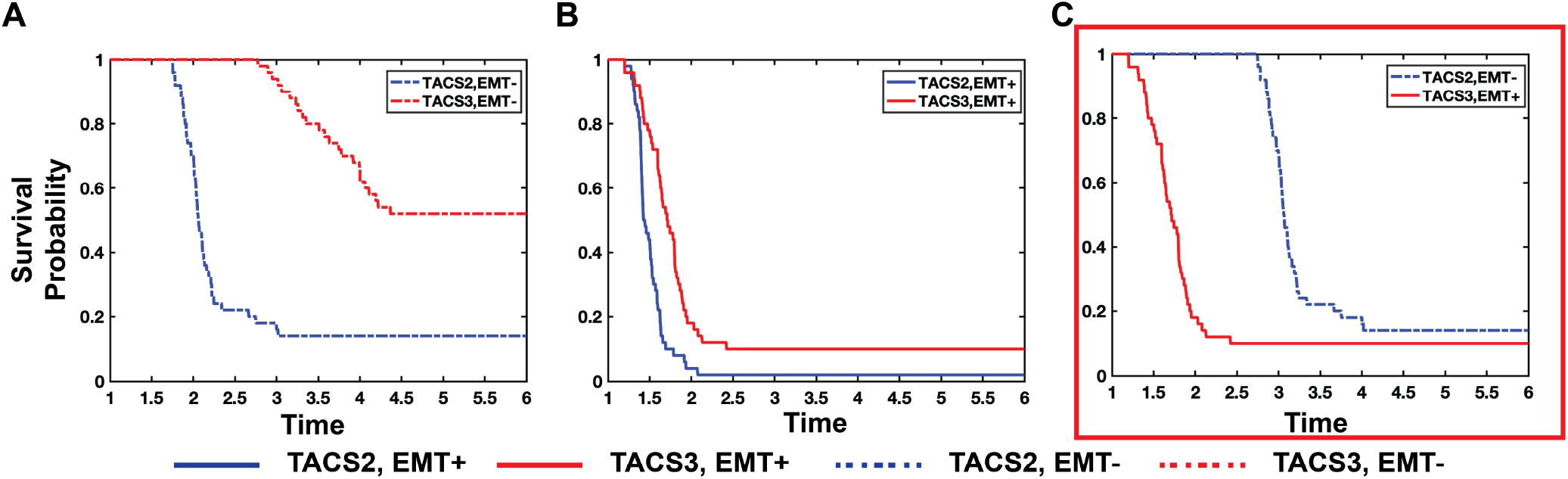
The concurrence of TACS3 and tumor adaptive changes is a significant factor contributing to the lower survival rates in TACS3 cases. The survival probability from 50 repeated experiments with varying TCR diversity in TACS2/EMT-, TACS3/EMT-(A), TACS2/EMT+, TACS3/EMT+ (B), TACS2/EMT-, TACS3/EMT+ (C). In all simulations, we assumed that clinical death occurs when the tumor size reaches ∼0.5cm. To simplify, we reduced the ROI size from a radius of 0.4cm to 0.25cm, meaning that the initial distance of T cells from the tumor is closer. λ=0.4, α_*t*_=0.01, α_*T*_ =1.6, *μ* =8 *·*10^−4^, *R*_*n*_=500∼2500, *r*_*n*_=100∼500.

To better understand the respective impacts of TACS3 and EMT on disease progression, we varied the occurrence of EMT and the timing of TACS3 initiation. We first modified the timing of TACS3 occurrence, which from our earlier findings was expected to prolong T cell residence times and infiltration efficiency (Figure 6D-E). We observed significant reductions in tumor burden with early TACS3, along with an earlier occurrence of substantial tumor killing (Figure 6A-B). While late TACS3 exhibits a higher peak in tumor burden, resulting in greater T cell expansion compared to early TACS3 (Figure 6B), the earlier onset of tumor killing in early TACS3 limits the time available for tumor evolution. These observations suggest that: 1) TACS3 facilitates enhanced T cell infiltration, which is consistent with our previous findings, 2) curtails the window for tumor evolution, and 3) partially alleviates challenges associated with tumor eradication. Hence, by controlling the timing of TACS3 emergence, we did not observe significant challenges to the immune system. Even when TACS3 appeared later, the efficient infiltration facilitated by TACS3 could assist T cells in encountering cognate antigens (Figure 6D).

**Fig. 6.**
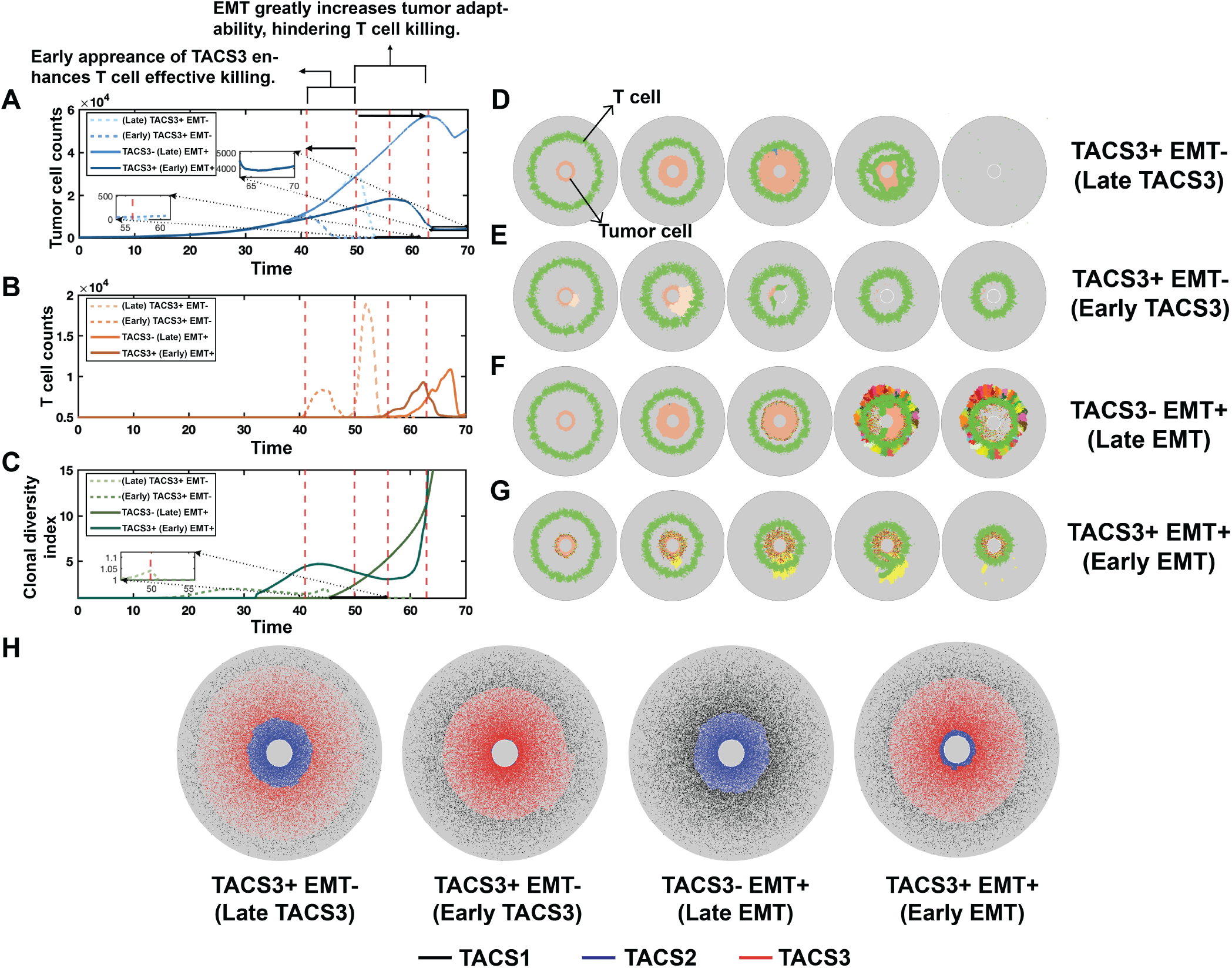
Tumor and T Cell interaction dynamics in the presence or absence of TACS3 and EMT. A: Tumor burden is depicted over time. The red dashed lines represent the time at which T cells initiate large-scale tumor killing in each condition. B: T cell counts are depicted over time. C: Clonal diversity index is depicted over time. Please refer to Methods for further details on the clonal diversity index. D-G: Snapshots of each condition at different times respectively. T cells are represented by green, while the remaining cells represent tumor cells, with different colors indicating distinct tumor clones. Among them, D: TACS3 occurs without EMT, and TACS3 occurs early. E: TACS3 occurs without EMT, and TACS3 occurs late. F: EMT occurs without TACS3. G: TACS3 and EMT occur simultaneously. H: Final distribution of fibers in each condition. TACS1 fibers are marked with black, TACS2 fibers are marked with blue, and TACS3 fibers are marked with red. In all conditions, λ=0.1, α_*t*_=0.02, α_*T*_ =1.8, *R*_*n*_ = 210, *r*_*n*_ = 10.

Given that TACS3 alone cannot fully explain the clinically observed survival in breast cancer (Figure 5A), we next assessed the impact of EMT occurring alongside TACS3. We identified three differences in EMT^−^ (Figure 6D-E) vs. EMT^+^ (Figure 6F-G) cases: Firstly, tumor burden is notably higher in EMT^+^ cases. Secondly, the time at which tumor burden begins to significantly decrease is later in EMT^+^ cases. Thirdly, there is a less dramatic decrease in tumor burden in EMT^+^ cases. These observations can be explained by higher levels of TAA-specific heterogeneity, reduced immunogenicity, and enhanced migration in EMT^+^ cases, all of which impair T cell killing. TACS3 enhances T cell infiltration relative to TACS2 cases (Figure 6A, F), thereby leading to earlier encounters between T cells and tumors and reductions in tumor burden (Figure 6F-G).

Collectively, these results suggest that phenotypic adaptation occurring late in TACS3 poses significant challenges to the immune system and reduces cancer elimination rates. From this, we conclude that the occurrence of TACS3 alone in our model cannot fully account for observed survival rates among breast cancer patients (15). Factors conducive to tumor fitness and adaptability concurrent with TACS3, such as EMT, are necessary to explain observed survival trends, and our model suggests that cancer phenotypic adaptive mechanisms significantly contribute to diminished survival outcomes (Figure S6B-C). This further substantiates our earlier hypothesis regarding the concurrence of EMT and TACS3. Under this assumption, we also compared the tumor evolution trajectories between TACS3^*−*^ and TACS3^+^ conditions with varying TCR recognition abilities; representative results are shown in Figure S7.

#### Increased checkpoint expression in TACS3 further elevates tumor escape rates

Given the impact of TACS and phenotypic adaptation on altered immunogenicity and subsequent expected cancer escape, along with the ability of mesenchymal tumor cells to upregulate PD-1 (Programmed Cell Death Protein-1)/PD-L1 (Programmed Death Ligand-1) expression by modulating certain pathways (43), we further explored how TACS-specific tumor evolution affects responsiveness to PD-1/PD-L1 inhibitor and its impact on patient survival. We next considered an increase in PD-1/PD-L1 levels at a certain time point, hindering T cell recognition of tumor cells. Based on our prior results, we simulated conditional combinations considered in Figure 5C. In our model, immune checkpoint up-regulation raises the threshold for T cell recognition of TAA. While there are multiple ways to account for this, we decided to allow cancer cells to become eliminated and recognized only if at least two TAAs were recognized since this uniformly impairs all T cell recognition in the presence of immune checkpoint. After a specific duration, we administered inhibitors to restore TCR recognition capability to its pre-elevated state. We conducted 10 replicates in each of the two settings, keeping all other parameters consistent. Representative images are depicted in Figure 7A,C.

**Fig. 7.**
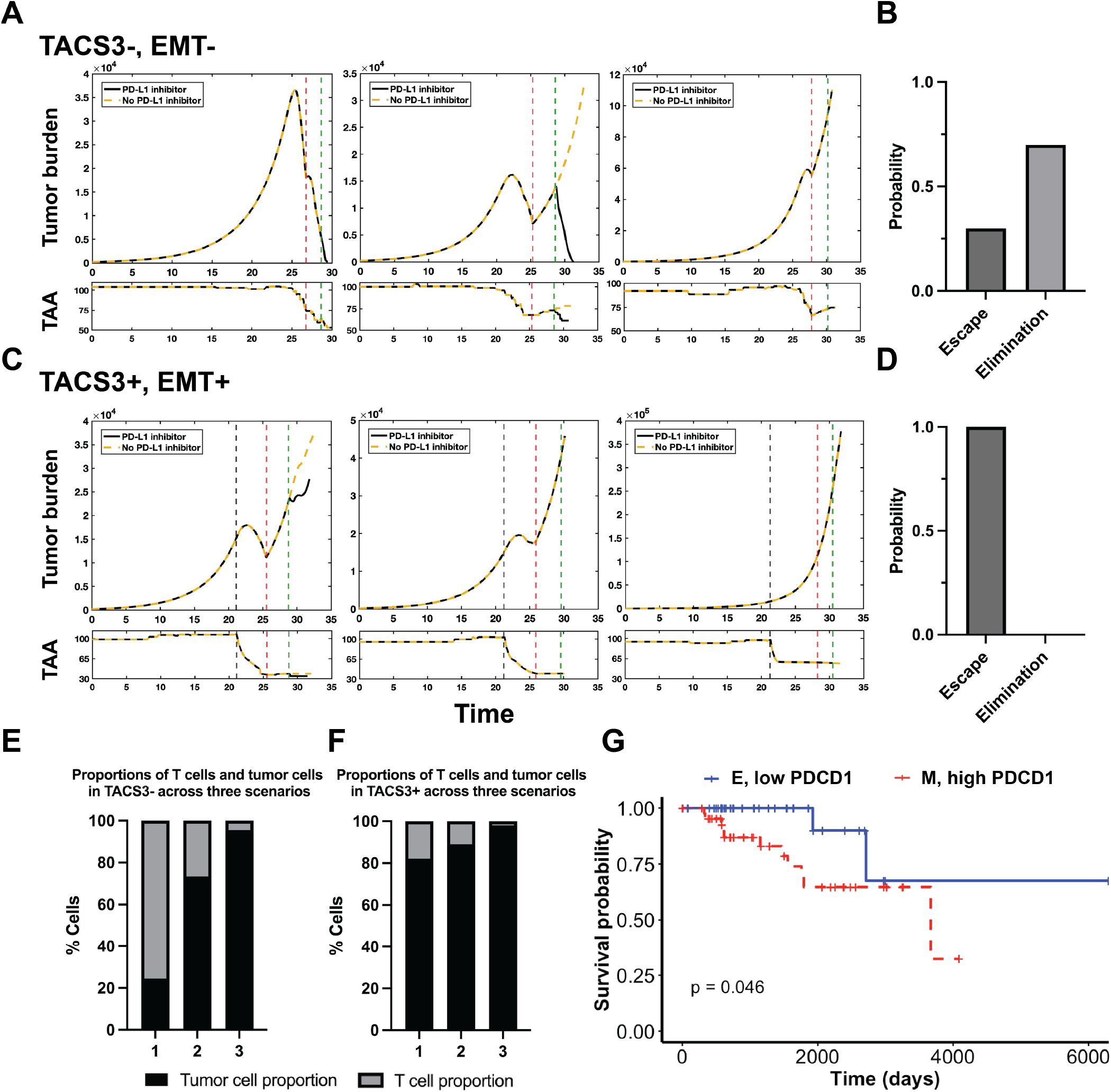
TACS-specific tumor evolution affects checkpoint inhibitor efficiency. A: Representative tumor burden dynamics and associated mean TAA counts in the TACS3-, EMT-conditions are depicted using identical model parameters. The red and green dashed lines represent the time of PD-L1 elevation and the administration of the PD-1/PD-L1 inhibitor, respectively. Three representative images are shown from left to right, illustrating three scenarios: (1) inhibitor responder, where tumors are eliminated regardless of inhibitor administration; (2) inhibitor responder, where inhibitor administration leads to tumor elimination, while without inhibitor, tumor escape occurs; (3) inhibitor non-responder, where tumors escape both with and without inhibitor administration. B: In the TACS3-, EMT-condition, the probability distributions of tumor elimination and escape after the addition of the inhibitor are depicted. C: Under the same parameter regime, the representative tumor burden dynamics and associated mean TAA counts in the TACS3+, EMT+ conditions are depicted. The gray dashed line represents the time of EMT occurrence. Two red dashed lines represent the time of PD-L1 elevation and the administration of the PD-1/PD-L1 inhibitor, respectively. In all three scenarios, tumors all escape. D: In the TACS3+, EMT+ condition, the probability distributions of tumor elimination and escape after the addition of the inhibitor are depicted. E, F: In both aforementioned settings, the proportions of tumor and T cells at the pre-treatment time point are depicted. G: The survival probability of two cohorts, E with low PDCD1 expression and M with high PDCD1 expression, in the TCGA BRCA database were analyzed. Please refer to Methods for more details. In all conditions, λ=0.2, α_*t*_=0.01, *α*_*T*_ =1.8, *R*_*n*_=500, *r*_*n*_=100.

In our repeated experiments, TACS3^*−*^ yielded a high variability of responders (leading to tumor elimination) and non-responders (indicating tumor escape; Figure 7A-B). Through an examination of the antigenic level of tumor clones, we found differing levels of immunoediting. Conversely, we consistently observed tumor escape in the TACS3^+^ setting, indicating either non-responsiveness or a lack of highly effective responses (Figure 7C-D). Here, EMT reduces the effective number of TAAs available for T cell targeting. Elevated PD-L1 levels further exacerbate this effect, presenting a considerable obstacle to the immune response. In the presence of a substantial tumor burden, even with the addition of inhibitors, there is minimal, if any, enhancement in cancer cell recognition. We further compared the proportions of T cells before the application of PD-L1 inhibition, and found that a higher proportion of T cells correlates with lower tumor burdens and favorable ultimate cancer elimination rates (Figure 7E-F). This observation aligns with recent findings regarding triple-negative breast cancer (TNBC) that the proliferative fractions of CD8+ T cells as the second most significant predictor of immune checkpoint blockade response, following MHC I&II. (44). Therefore, we posit that the concomitant occurrence of EMT and immune checkpoint together in TACS3 significantly diminishes T cell recognition and hence immune-mediated cancer elimination, which consequently reduces the responsiveness of checkpoint inhibitors.

To validate our findings, we conducted a survival analysis on breast cancer (BRCA) patients from The Cancer Genome Atlas (TCGA) database. We utilized E-cadherin (CDH1) and Vimentin (VIM) as markers to delineate epithelial (E) and mesenchymal (M) cohorts, respectively (45). Each cohort comprised 40 genes associated with its respective marker, constituting the gene signatures for the E and M cohorts (Figure S9A-B; please refer to the Methods for full details) (46–48). We compared survival differences between the two cohorts and although general trends were in agreement with our model predictions, we found no statistical significance (Figure S9C). We observed similar findings when considering PDCD1 up-regulated vs. down-regulated cases (Figure S9D-E). However, when patients were simultaneously stratified according to EMT and PDCD1 status, we identified a statistically significant reduction in overall survival for patients with cancer cells having M/PDCD1+ signatures, relative to those with E/PDCD1-signatures (Figure 7G). These findings are in agreement with our model predictions and offer an explanation for how phenotypic adaptation and immune checkpoint together can impair T cell recognition.

## Discussion

In solid tumors, ECM topology is known to be an important feature of the TME that affects tumor progression, metastasis, and therapeutic resistance (49, 50). However, the precise way in which cancer growth and tumor-immune coevolution are affected by this topology remains unclear. To begin to address this, we developed EVO-ACT model to explore the dynamic and spatial aspects of immune cell and tumor interaction as a function of TACS. We applied our model to gain quantitative insights into several dynamic features of this complex process, including how TACS affects both tumor cell and T cell migration efficiency. Our findings suggest that TACS exert a greater impact on chemokine-driven T cell infiltration, and they do not constitute an absolute barrier to T cell infiltration. Our findings predict that TACS-specific T cell infiltration patterns influence immunoediting, and TACS3, together with late-stage phenotypic adaptations like EMT and elevated PD-L1 expression, collectively contribute to reduced predicted patient survival and decreased responsiveness to PD-L1 inhibitors.

Our work identified the impact of TACS in cell migration direction, spatial distribution and quantified the cell invasion efficiency in three common TACS, with TACS3 *>* TACS1 *>* TACS2. The highest migration efficiency observed in TACS3 offers new insights into the enhanced cell migration observed in TACS3 compared to random fiber environments (34). Another widely accepted explanation posits that tumor cells in TACS3 exhibit fewer protrusions, resulting in increased directional persistence and, consequently, enhanced invasion efficiency (35). Additionally, our findings indicate that TACS does not constitute an absolute barrier to cell invasion, which has been debated in recent years (13, 14, 16, 17); rather, it influences cell invasion efficiency significantly. However, the variation in invasion efficiency induced by TACS differs between tumor cells and T cells. Under the directed influence of chemokines, T cells are more profoundly affected by TACS. This finding also suggests that adopting stroma-mediating treatments may offer greater assistance to T cells. Prior work has underscored the growing recognition of the importance of ECM geometry in influencing cell invasion and deepening our understanding of the elevated invasion efficiency in aligned fibers (22, 34, 51, 52). Future studies should build upon this foundation by considering that modifying the TACS environment may have differing impacts on tumor cells and T cells.

Since TACS does not completely hinder T cell infiltration, our analysis focused on highlighting the heterogeneous T cell spatial distributions in each TACS. Prior work has illustrated the importance of including chemokine attraction and antigen specificity for generating observed T cell spatial distributions (32). Our results demonstrate that TACS also plays a significant additional role in generating observed T cell spatial distributions.Given the significant role of TACS in allocating spatial positions for tumor clones, we argue that the spatial distribution of T cell clones is also influenced by TACS, which in turn influences the location of expanding T cell clones (Figure S5). An alternative theoretical approach focused on T cell distributions in the tumor core, invasive margin, and tumor stroma. Significant differences in T cell distribution and function have been observed in these three distinct regions, with notable characteristics identified in each. T cells within the tumor core exhibit tight interactions with tumor cells due to their close proximity (53). The prolonged stimulation from tumor antigens may be a contributing factor to T cell exhaustion, supported by recent research demonstrating that chronic tumor antigen/TCR stimulation reinforces epigenetic programs associated with dysfunctional hallmarks, resulting in dysfunctional T cells (32, 54). These results are consistent with prior findings that T cell dysfunction and exhaustion correspond to higher expression of co-stimulatory molecules such as PD-1, TIM-3, and VISTA (55, 56). This phenomena is comparable to our model (Figure S5) in settings where T cells that have interacted with the tumor are assumed to be exhausted, leading to the concentration of potentially exhausted T cells inside the tumor. In contrast, T cells in the invasive border are believed to have higher density, functionality, and a weak expression of co-inhibitor molecules compared to those in the tumor core or stroma (55, 57, 58). In our model, T cells migrate toward the tumor, at which point factors such as antigen specificity, inflammatory cytokines, and distinct TACS lead to varying distributions of expanding T cells within the tumor. However, we found that the majority of T cell clones are still predominantly located at the tumor margin.

Our results demonstrate that TACS modulate T cell infiltration efficiency, which in turn influences tumor evolution through distinct levels of immunoediting. TACS-specific changes in T cell infiltration therefore result in varying degrees of selective pressure on tumors. Under conditions of heightened T cell infiltration and recognition, potent negative selective pressure drives surviving cancer cells to have reduced immunogenicity (2). This heterogeneity of T cell infiltration and neoantigen-derived immunoediting has been observed in various cancers, such as lung cancer and breast cancer (2–4, 17), which can be achieved through loss of heterozygosity in human leukocyte antigens, depletion of expressed neoantigens or decreased TAAs presentation efficiency (2, 4). Furthermore, studies indicate that the distribution of tumor mutation burden (TMB) is heterogeneous, and tumors with heterogeneous infiltration are more prone to manifesting a heterogeneous TMB (2, 26, 59). Our results corroborate this finding, as demonstrated in Figure S10. We propose that TACSs also plays a significant role in inducing TMB heterogeneity.

While we have shown that TACS may influence tumor evolution in a variety of ways, our results indicate that one potential reason for the lower survival rates observed in TACS3^+^ cancer patients is the concomitant occurrence of phenotypic adaptation (15). In particular, clinically observed survival trends in TACS3^+^ and TACS3^*−*^ cases in breast cancer were only explained in our modeling framework by incorporating additional phenotypic adaptation mechanisms, including EMT (15). Our results suggest that TACS3 may not be the primary driver of lower survival rates. EMT provides one possible explanation wherein heterogeneous and invasive clones are facilitated in a TACS3^+^ environment to metastasize (12, 30, 34, 52, 60, 61). Other relevant phenotypic adaptation mechanisms may also actively affect these results, and such distinct mechanisms may occur and be of direct importance for distinct cancer subtypes. Relevant mechanisms for metastatic disease are marked by higher growth rates and lower antigenicity compared to their ancestors, making them more resistant to elimination (62). Moreover, we propose that such cellular phenotypic alterations typically manifest in the late stages. Otherwise, if occurring early, such as during the TACS2 phase, the rapid escalation of tumor heterogeneity and the hindrance posed by TACS2 on T cell infiltration would expedite tumor evasion rapidly (Figure 6F). Our results further indicate that this TACS-specific tumor evolution trajectory also influences the efficacy of checkpoint inhibitors. The different degrees of immunoediting caused by TACS affect the responsiveness of PD-L1 inhibitors (44). Our findings also indirectly suggest that the earlier use of inhibitors may lead to better outcomes, underscoring the importance of early differentiation between inhibitor responders and non-responders.

Our foundational model makes a number of assumptions. Firstly, in our model, we assume that all cells at the tumor boundary undergo EMT simultaneously upon reaching the EMT threshold. Also, we assume that the mutation rates of all tumor clones are identical. However, in reality, different tumor clones may undergo EMT and mutation at different times and with different rates, potentially leading to larger variations in growth, migration, and collective migration across distinct intratumoral subpopulations. Secondly, our model does not account for the presence of exhausted T cells; T cells are either in an activated state or have died and are subsequently removed from the system. We expect that in reality exhausted T cells that persist within the tumor core provide additional hindrance to T cell killing. It is also likely that the pro- vs. anti-tumor characteristics of the immune microenvironment, which we did not model in detail here, further determine the extent to which this exhaustion occurs. The presence of exhausted T cells and their occupation of space, along with increased metabolic demands such as oxygen consumption, further exacerbates the difficulty of T cell killing in real-life situations compared to our model. Thirdly, our framework models tumor growth and tumor-immune interactions in a 2D space, whereas real tumor populations develop in three dimensions. Our analysis considered idealized TACS2 and TACS3 collagen arrangements for simulations when in reality a number of variable and overlapping topologies likely exist. The incorporation of patient-specific ECM orientation is an important next step and the focus of future research efforts. Lastly, the current model only considers the interaction between tumor cells and T cells, yet many other features in the TME affect T cell recognition, including dendritic cells, tumor-associated macrophages (TAM), and metabolic and chemical signatures. Nonetheless, our foundational model is able to provide a population dynamical explanation for a variety of relevant characteristics of the tumor-immune interaction.

Taken together, our results quantify the impact of ECM architecture on the tumor-immune interaction, and our modeling approach represents a novel mathematical framework to incorporate collagen-specific information in to a description of tumor and immune co-evolution. We also speculate that TACS may affect all elements in the TME reliant on fiber movement or requiring ECM penetration, potentially influencing tumor evolution or treatment outcomes. Hence, our findings also indirectly affirm the importance and necessity of adopting stroma-modifying treatments in clinical practice. Moreover, since ECM architecture does not exist independently, further research is required to investigate the collective effects of various TACS byproducts, such as angiogenesis, spatial distribution, interactions of different cells, and varying microenvironments, on the tumor-immune interaction.

## Methods

### Initial conditions and model structure

The implemented program utilizes Gillespie’s simulation algorithm to model stochastic events for two agents: tumor cells and T cells. Tumor cell events include invasion, division, and migration, each governed by rates *γ, λ, α*_*t*_ respectively. Successfully divided cells undergo mutation based on the mutation rate, *μ* per cell division. Initially, tumor cells are assumed homogeneous. T cells only perform movement, with a rate *α*_*T*_. We define an ROI with radius *R*, containing a circular tumor mass centrally with radius *r*. Tumor cells inside this central circle (*n ∼* 4000) are not tracked until they invade beyond it. Upon invasion, cells can migrate or divide. Initially, we assume that 200 tumor cells have invaded the central mass. T cells (*N* = 5000) are initially positioned at the ROI boundary and commence infiltration towards the tumor, as illustrated in Figure S1. Some important parameters are listed in Table S1.

### TACS generation and remodeling

Collagen fibers occupy the annular space between the central tumor and the surrounding ROI. Fiber density is imposed along a linear gradient maximal at the tumor boundary. (12). The lengths of these fibers follow a normal distribution (mean *μ* = 10*μ*m; variance *σ*^2^ = *μ/*10). Initially, all fibers are randomly packed, and their directions are normally distributed, forming TACS1. Remodeling is accounted for by allowing cancer cells at the tumor boundary to change the fiber orientation. With each tumor cell division, fibers within a radius of *r*_2_ = 75*μ*m are remodeled into perfectly aligned TACS2. The remodeled fibers will orient perpendicular to the line linking the dividing cell’s center and the center of the fiber undergoing remodeling. During EMT concurrent with TACS3, cells on the tumor periphery remodel fibers within a radius of *r*_3_ ∼ 0.18*cm* into perfectly aligned TACS3. This remodeling is based on previous findings suggesting that tumors can alter fibers within a range of 5 times the original tumor spheroid diameter into TACS3 (25). We also assume that once fibers are remodeled into TACS3, they cannot revert to TACS2. In the article, we employ the terms TACS2 and TACS3 to represent the consistent distribution of fibers throughout the entire ROI as either TACS2 or TACS3. We also assume higher alignment closer to the tumor center and lower alignment farther away (12). This is equivalent to the tumor no longer undergoing remodeling of fibers under a deterministic fiber architecture. Conversely, we use TACS3^−^ and TACS3^+^ to denote the transition of TACS from TACS1 to TACS2 or TACS3, indicating that the tumor will gradually remodel fibers with the occurrence of division or EMT.

### Tumor division

We consider population dynamics for individual clones in the population. Specifically, When tumor division occurs, we first calculate the total rate of division for each tumor clone based on its size and division rate. Subsequently, we uniformly select a cell at random for division from any tumor clone according to its total division rate, ensuring that the chosen position for division does not overlap with any existing tumor cell. During each division time window, we allow a maximum of 100 cells to attempt division, with each cell having up to 200 attempts to select a division position. Upon successful division, the newly divided cell remodels fibers within a range of *r*_2_ from TACS1 to perfectly aligned TACS2. If no suitable division position is found for a cell, no division occurs. Following division, each newly divided cell undergoes mutation based on the mutation rate, *μ*.

### Tumor mutation

We assume that each tumor clone possesses a certain quantity of TAAs, following a Poisson distribution with a mean of n = 100. Mutated cells randomly adjust the quantity of antigens, either increasing or decreasing. The variance in adjusted antigen levels also follows a Poisson distribution with a mean of 10. The division rate of the mutated tumor clone will increase or decrease by 1 % of *λ* with each gain or loss of a TAA. Conversely, if no mutation occurs, the new tumor cell retains the same properties as the original tumor cell, including division rate, migration rate, and antigen set.

### Tumor migration

When tumor migration occurs, we compute the migration probability for each tumor clone, akin to calculating the division probability. Subsequently, we randomly select a tumor clone based on its migration probability and then randomly choose a cell from within that clone for migration. Each tumor cell randomly selects a fiber within a range of *ϵ*_*t*_ = 75*μ*m as the migration direction. There’s an equal probability of choosing either the selected fiber’s direction or its opposite direction. The distance traveled by the tumor cell is determined by the diffusion coefficient D and the current time window (t). Each step in the EVO-ACT model checks for issues related to spatial overlap to ensure that neither the migration path nor the destination of each tumor cell overlaps with any other tumor cell.

### EMT

In our model, the initiation of EMT depends on the tumor burden. Once the cumulative tumor burden exceeds a specified threshold, which varies in each condition, all divisible tumor cells undergo EMT simultaneously. EMT induces changes including a fivefold decrease in tumor division rateλ, a fivefold increase in migration rate *α*, and a reduction in tumor clone immunogenicity, resulting in a random decrease in tumor antigen quantity following a Poisson distribution with a mean of 15. Moreover, all cells undergoing EMT will remodel fibers within a range of *r*_3_ to TACS3.

### T cell migration, killing, and expansion

When T cell migration occurs, we randomly select a T cell for migration. The direction of T cell migration is influenced by both the gradient of tumor-secreted chemokines and the orientation of surrounding fibers within a range of *ϵ*_*T*_. The migration distance of the T cell depends on the intensity of the chemokine gradient (ΔC), the local fiber density, and the duration of the current time window (t). We ensure that neither the migration path nor the destination of the T cell overlaps with any other T cell. To model the specificity of individual TCRs, we assume that each T cell can recognize only one type of antigen. If migration is successful, the T cell checks all tumor cells within a range of *ϵ*_*k*_ = 80*μ*m for a recognizable antigen. The number of checking attempts is proportional to the duration of the current time window (t). If a recognizable antigen is found within *‘*_*k*_, the T cell initiates killing. The killed tumor cell is removed from the system, and the T cell executing the killing generates a new T cell to occupy the position of the dead tumor cell. The newly generated T cell inherits the same TCR as its parent T cell. However, unlike its parent cell, all newly generated T cells have a survival window, whereas those T cells present in the model from the beginning do not possess such a survival window. Although memory cells are not explicitly incorporated into the model, the above dynamics create an effective memory of antigen-specific T cells while also accounting for the dynamics of T cell expansions and contractions. We assume that all T cells with survival windows have equal survival periods, and their timers start upon birth. Once their designated survival period elapses, they are assumed to have died and are removed from the system, as depicted in Figure S2.

### Clonal diversity index

To measure clonal diversity, we used the inverse Simpson index defined as *D* = 1*/ Σ* _*i*_ (*p*_*i*_)^2^, where *p*_*i*_ is the frequency of the ith combination of driver mutations.

### Identification of EMT-associated gene signature

In the TCGA BRCA database, we selected 610 cases and identified the 40 genes most correlated with E-cadherin (CDH1) and Vimentin (VIM) using Spearman correlation analysis (45). These genes were chosen as the gene signatures for the E and M groups. Building upon previous research indicating the presence of hybrid phenotypes within E and M categories (46), we defined the E and M cohorts based on the expression levels of the gene signatures in the selected 610 cases (Figure S9A-B). The cutoff values were determined by Equation 1,2. We selected samples with simultaneous high expression of E gene signature and low expression of M gene signature as the E cohort. Conversely, samples with simultaneous low expression of E gene signature and high expression of M gene signature were chosen as the M cohort. This was done to minimize the risk of selecting E and M signatures that contained hybrid E/M intermediate phenotypes (46–48).

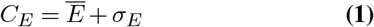

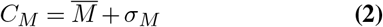

where *C*_*E*_ and *C*_*M*_ are the cutoff values of E and M cohorts respectively, *Ē* and 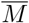 are the mean expressions of E and M gene signatures respectively, σ_*E*_, and σ_*E*_ are the standard deviations of E and M gene signature expressions respectively.

### Identification of PDCD1 high and low expression cohorts

After distinguishing the E and M cohorts, we calculate the median of PDCD1 gene expression in these two cohorts as the cutoff value (Figure S9D). Subsequently, we further divide the E and M cohorts into high and low groups based on PDCD1 expression.

## Acknowledgment

We express our gratitude to Dr. Feng Zhao, Alvis Chiu for providing fruitful discussions on this study. JTG was supported by the Cancer Prevention Research Institute of Texas (RR210080). JTG is a CPRIT Scholar in Cancer Research.

## Data availability

The code and part of the raw data for the EVO-ACT model are available on the following GitHub website: https://github.com/TAMUGeorgeGroup/EVO-ACT.

