## Supplementary Information for "Distinct evolutionary patterns of tumor immune escape and elimination determined by ECM architectures"

| Variables | Notations | Values | References |
|---|---|---|---|
| Radius of central tumor circle | r | $5 \times 10^{-2}$ cm | |
| Radius of ROI | R | 0.4 cm | |
| Tumor cell diameter | d | 15 $\mu$m | (63) |
| T cell diameter | D | 8 $\mu$m | (64) |
| Tumor division rate | $\lambda$ | 0.1 $\sim$ 2 | |
| Tumor migration rate | $\alpha_t$ | 0.01 $\sim$ 1 | |
| Tumor mutation rate | $\mu$ | $5 \times 10^{-4} \sim 8 \times 10^{-4}$ | |
| Number of tumor cells in the central tumor circle | n | 4444 | |
| Initial number of T cells | $N$ | 5000 | |
| T cell migration rate | $\alpha_T$ | 1.6 $\sim$ 1.8 | |
| Mean antigen counts in tumor clones | A | 100 | |
| Total number of fibers in ROI | $N_f$ | 8000 | |
| Mean fiber length | l | 10 $\mu$m | (23) |
| TACS2 remodeling radius | $r_2$ | 5d=75 $\mu$m | |
| TACS3 remodeling radius | $r_3$ | $\sim$120d=0.18 cm | |
| The radius for a tumor cell to randomly pick a nearby fiber for movement | $\epsilon_t$ | 5d=75 $\mu$m | |
| The radius for a T cell to randomly pick a nearby fiber for movement | $\epsilon_T$ | 10D=80 $\mu$m | |
| The radius for a T cell to randomly pick a nearby tumor to kill | $\epsilon_k$ | 3d=45 $\mu$m | |

**Table S1.** Values of key parameters in the model and associated references.

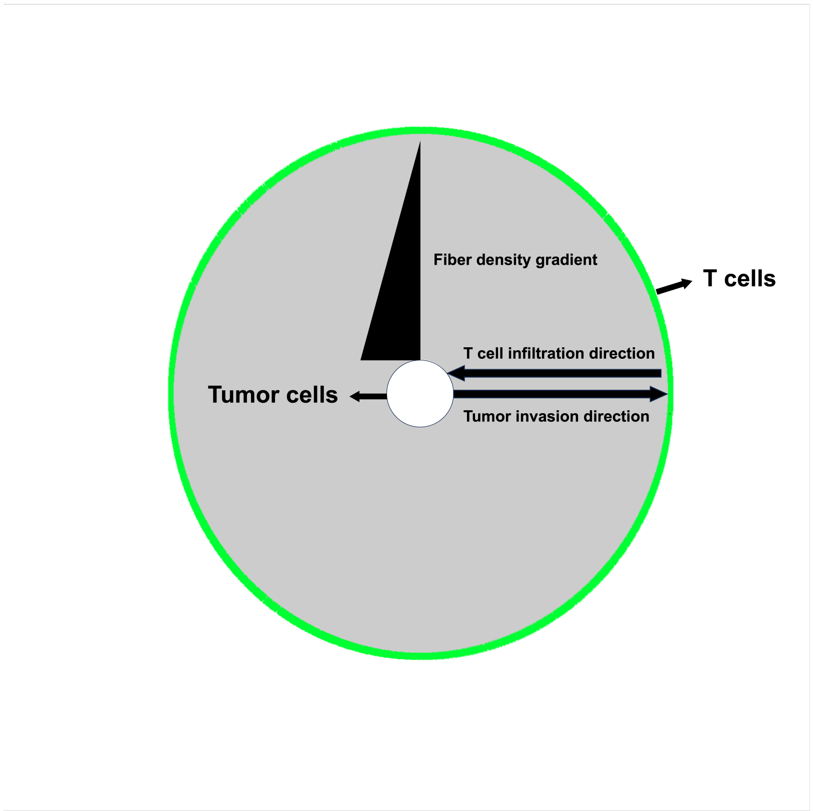


**Fig. S1. The initial spatial configuration of tumor Cells, T Cells, and fibers in our model.** The circular region represents our ROI, within which a circular tumor region is initially defined. Tumor cells within this circular area must invade beyond its boundaries to migrate or divide. Those that invade will migrate outward. All T cells are assumed to be positioned at the boundary of the ROI and will infiltrate towards the tumor. The gray area in the illustration denotes the presence of fibers. In the simulations to follow, we impose a decreasing fiber density that varies linearly as a function of the radius.

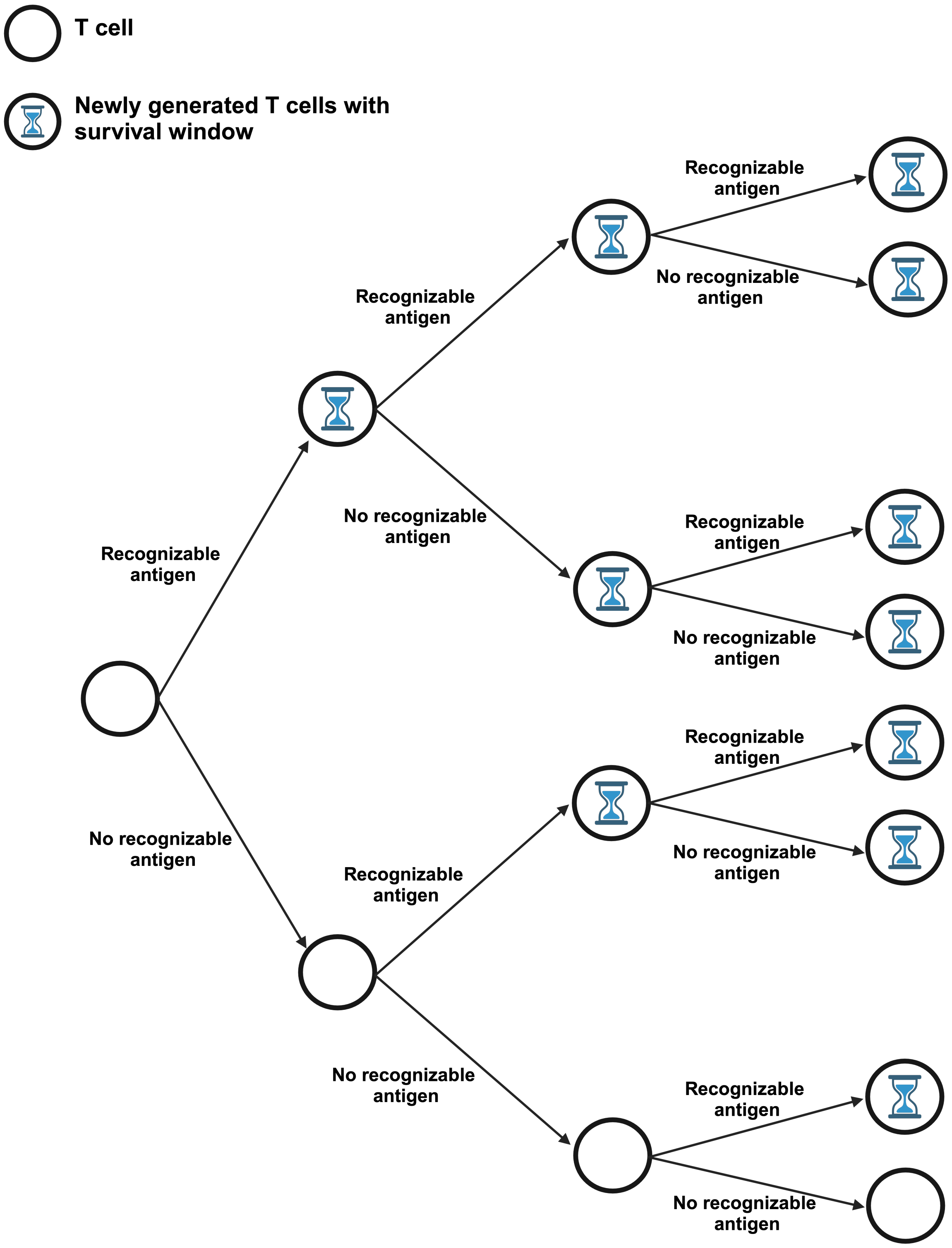


**Fig. S2. Rules for T Cell expansion.** In the designated ROI, existing T cells will expand upon encountering cognate tumor antigens. These newly generated T cells share identical lifespans, starting a countdown timer upon birth. Upon its expiration, these T cells are removed from tracking.

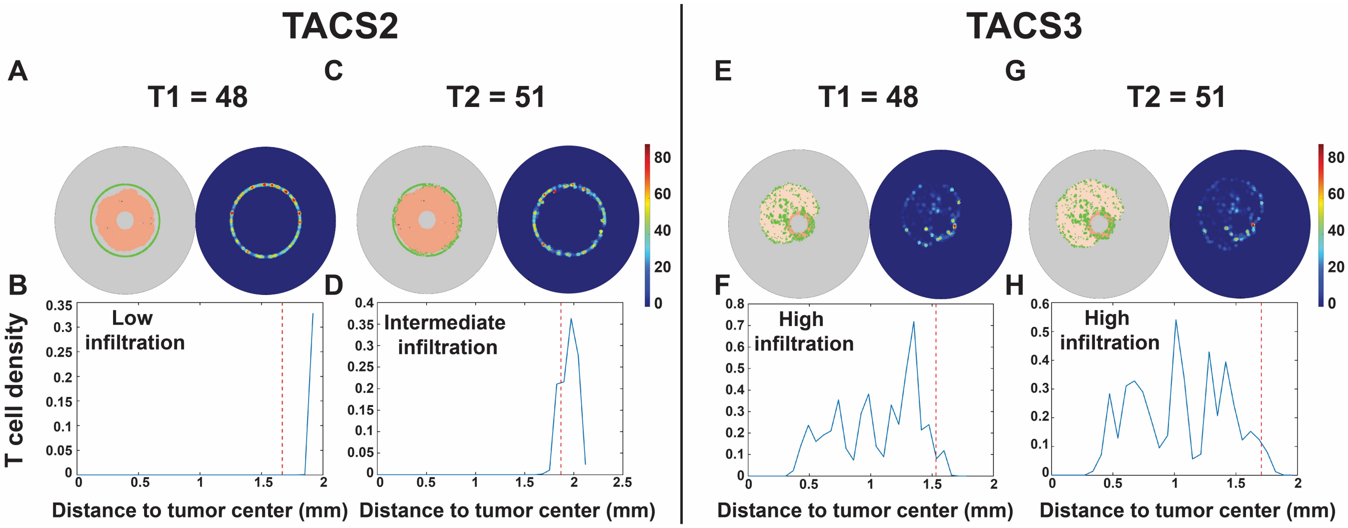


**Fig. S3. TACS affects the spatial distribution heterogeneity and infiltration level of T cells. TACS3 facilitates more efficient T cell infiltration.** In perfectly aligned TACS2 (A-D) and TACS3 (E-H), representative spatial distributions for T cell infiltration were quantified with respect to their distance to the tumor center at the same time point. The red dashed lines in the graph represent the boundary of the tumor core. For the same time duration, T cells in TACS3 can infiltrate into the interior of the tumor more efficiently.

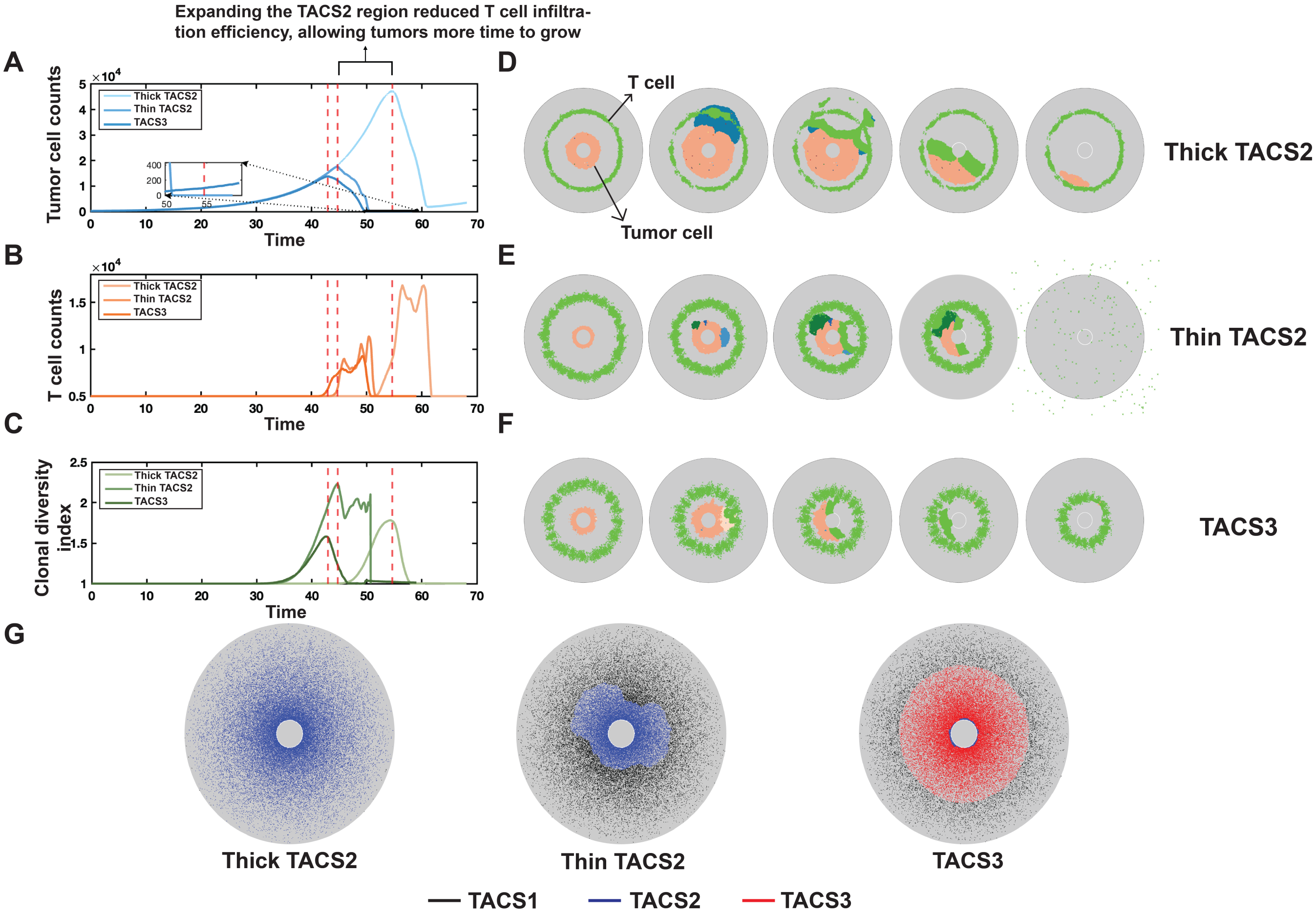


**Fig. S4. Tumor and T Cell interaction dynamics in varied TACS2 and TACS3 configurations.** A: Tumor burden is depicted over time. The red dashed lines represent the time at which T cells initiate large-scale tumor killing in each condition (defined by the time of maximal tumor size). B: T cell counts are depicted over time. C: Clonal diversity index is depicted over time (see Methods for full details). D-F: Snapshots of each condition at different times respectively. T cells are represented by green, while the remaining cells represent tumor cells, with different colors indicating distinct tumor clones. Among them, D: Thick TACS2 region. E: Thin TACS2 region. F: A comparison of TACS3 condition. G: Final distribution of fibers in each condition. TACS1 fibers are marked with black, TACS2 fibers are marked with blue, and TACS3 fibers are marked with red. In all simulations, $\lambda$=0.1, $\alpha_t$=0.02, $\alpha_T$=1.8.

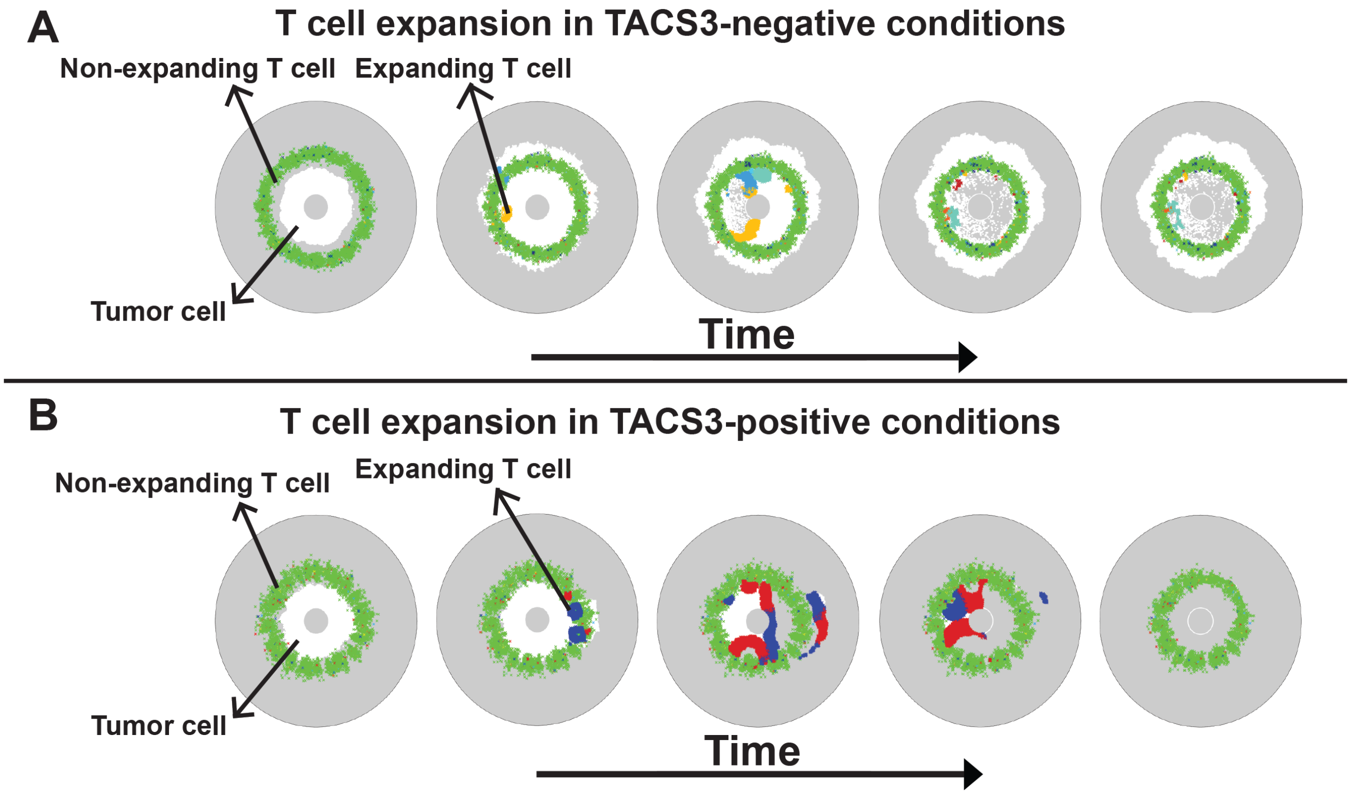


**Fig. S5. TACS further influences the spatial distribution of expanding T clones by influencing the spatial distribution of tumor clones.** The spatial distribution of T cell expansions against tumor clones are illustrated through time for representative cases involving A: TACS3$^{-}$ and B: TACS3$^{+}$ conditions. All tumor cells are marked in white. Non-expanding T clones are represented in green while expanding T clones are marked with colors excluding green ($\lambda$=0.1, $\alpha_t$=0.02, $\alpha_T$=1.8, $R_n$ = 210, $r_n$ = 10 in each case).

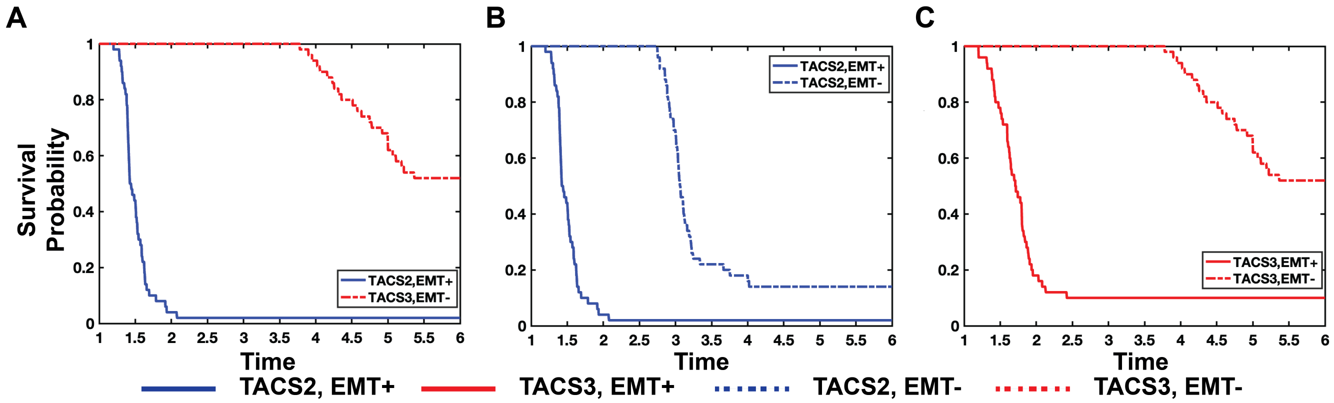


**Fig. S6. In our model, TACS alone cannot account for the observed low survival rates in TACS3$^{+}$ cases. Elevated cell adaptation significantly reduces survival rates in both TACS2 and TACS3.** The survival probability from 50 repeated experiments with varying TCR diversity in TACS2/EMT$^{+}$, TACS3/EMT$^{-}$ (A), TACS2/EMT$^{+}$, TACS2/EMT$^{-}$ (B), TACS3/EMT$^{+}$, TACS3/EMT$^{-}$ (C). In all simulations, we assumed that clinical death occurs when the tumor size reaches ~0.5cm. To simplify, we reduced the ROI size from a radius of 0.4cm to 0.25cm. $\lambda$=0.4, $\alpha_t$=0.01, $\alpha_T$=1.6, $\mu = 8 \cdot 10^{-4}$, $R_n$=500~2500, $r_n$=100~500.

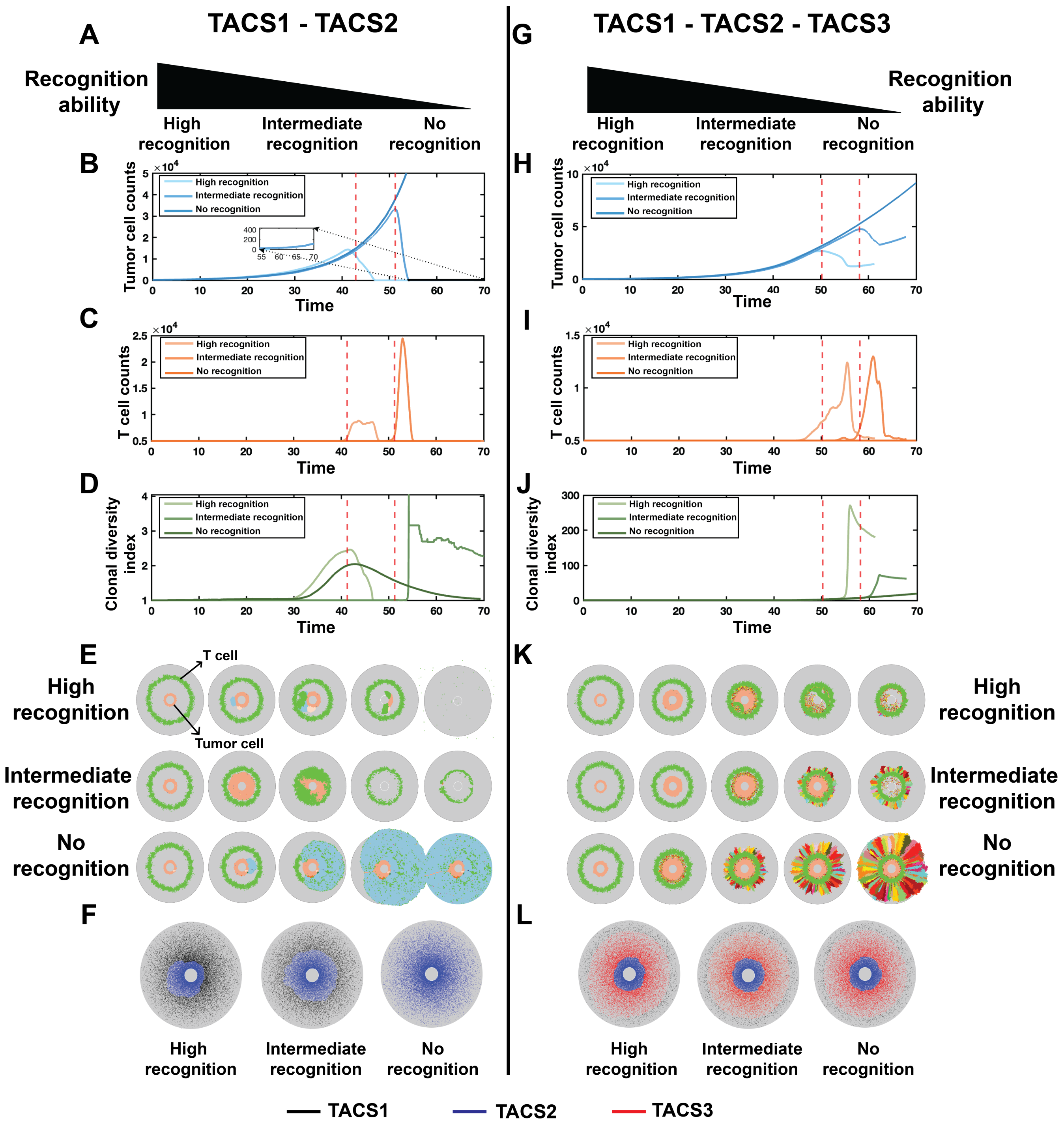


**Fig. S7. Tumor-T cell dynamics during TACS transitions.** A, G: In each of the three conditions for both TACS1 to TACS2 transitions and TACS1 to TACS3 transitions, it was assumed that the recognition ability of T cells gradually diminishes. B-D: Tumor burden, T cell counts, and clonal diversity index in three conditions (high recognition, intermediate recognition, and no recognition) during TACS1 to TACS2 transition are depicted respectively. Please refer to Methods for further details about the clonal diversity index. The red dashed lines represent the time at which T cells initiate large-scale tumor killing in each condition. In high recognition: $R_n$ = 210, $r_n$ = 210. In intermediate recognition: $R_n$ = 210, $r_n$ = 10. In no recognition: $R_n$ = 210, $r_n$ = 0. H-J: Tumor burden, T cell counts, and clonal diversity index in three conditions (high, intermediate, and no) during TACS1 to TACS2 to TACS3 transitions are depicted respectively. In high recognition: T cells can recognize all tumor clones that have not undergone EMT, and some clones that have undergone EMT. In intermediate recognition: T cells can recognize all tumor clones that have not undergone EMT, and cannot recognize any tumor clones that have undergone EMT. In no recognition: T cells cannot recognize any tumor clones. E, K: Snapshots of each condition at different times respectively. T cells are represented by green, while the remaining cells represent tumor cells, with different colors indicating distinct tumor clones. F, L: Final distribution of fibers in each condition. TACS1 fibers are marked with black, TACS2 fibers are marked with blue, and TACS3 fibers are marked with red. In all conditions, $\lambda$=0.1, $\alpha_t$=0.02, $\alpha_T$=1.8.

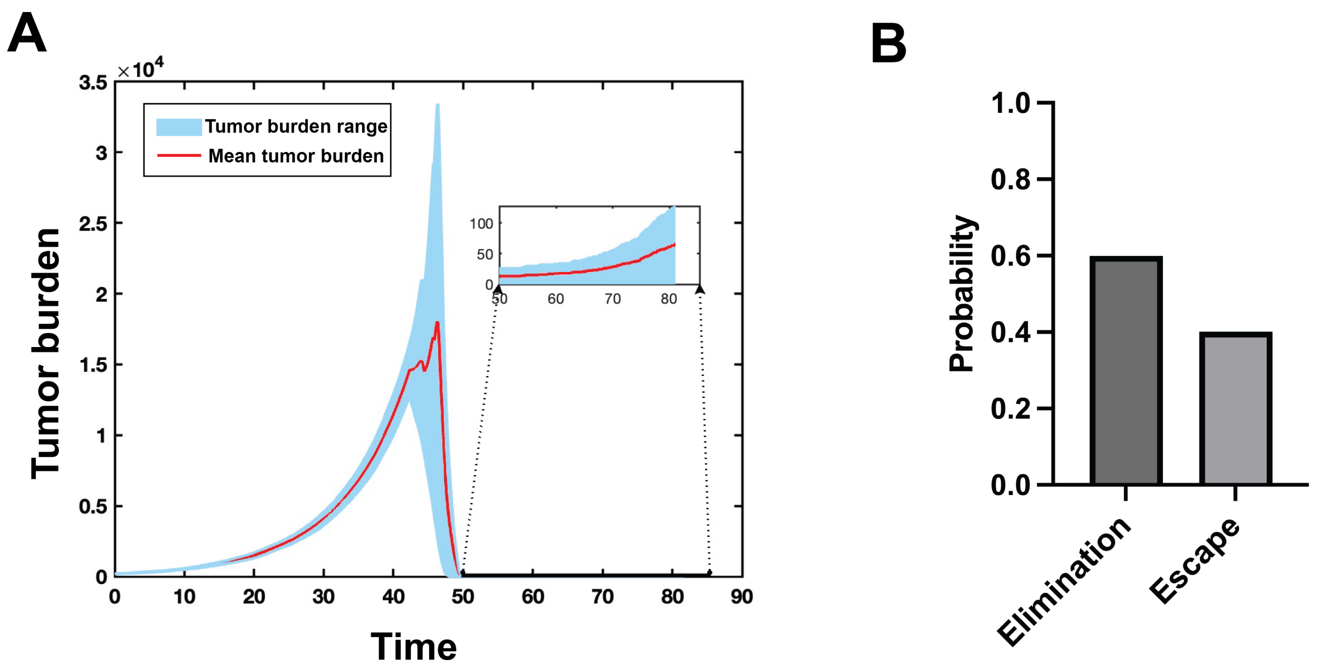


**Fig. S8. 15 repetitions of intermediate recognition during the transition from TACS1 to TACS2.** We conducted 15 repetitions of the intermediate recognition condition depicted in Figure S7E. A: Range and average value of tumor burden. B: Probability distribution of tumor evasion and elimination across the 15 repetitions.

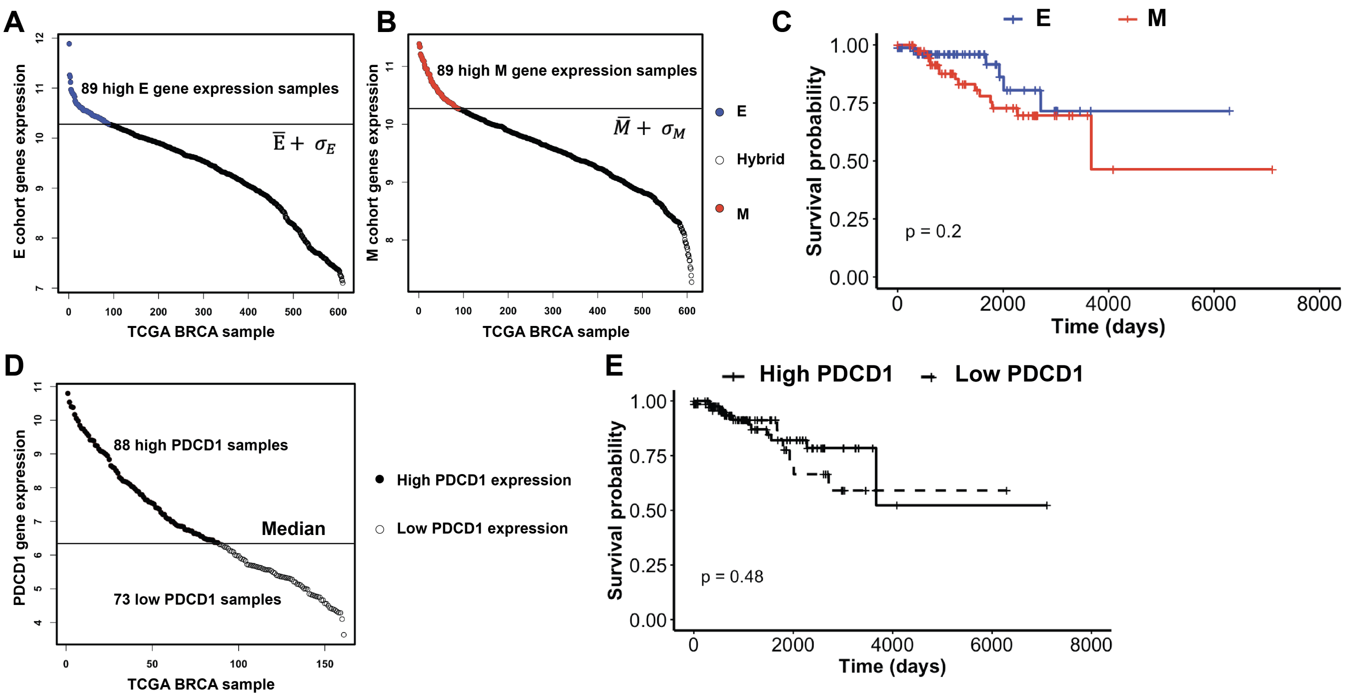


**Fig. S9. Identification of E and M cohorts and PDCD1 high and low expression cohorts.** A, B: In the selected 610 cases, we observed the expression status of E and M gene signatures. Based on the cutoff value, samples with expression levels above the cutoff value were selected for the E or M cohort. C: The survival probability of selected E and M cohorts, with p = 0.2. D: In the selected E and M cohorts, samples with PDCD1 expression above the median PDCD1 expression are categorized as the high PDCD1 expression group, while the remaining samples are classified as the low PDCD1 expression group. E: The survival probability of both high and low PDCD1 expression group, with p = 0.48.

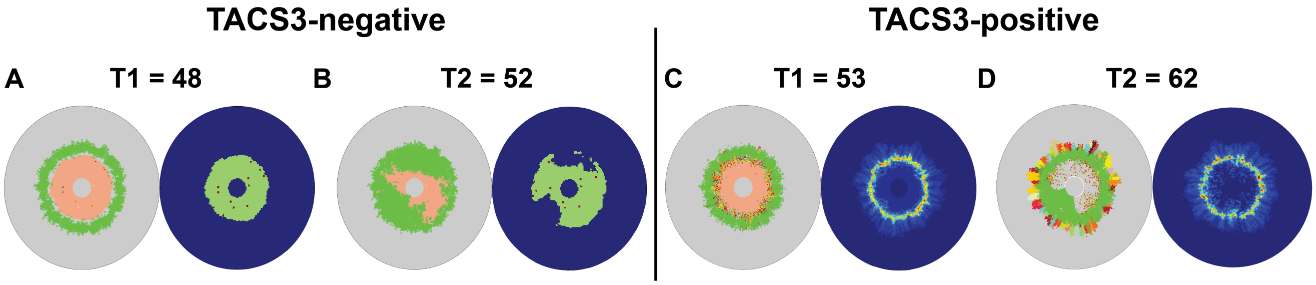


**Fig. S10. The heterogeneity of TMB between TACS3$^{-}$ and TACS3$^{+}$ conditions.** Under conditions identical to those in Figure 4A-H of the main text, we further quantified the spatial distribution of TMB based on the number of tumor clones surrounding each tumor. We posit that TACS further influences the spatial distribution heterogeneity of TMB by affecting tumor spatial distribution, which is consistent with previous findings (2, 26).

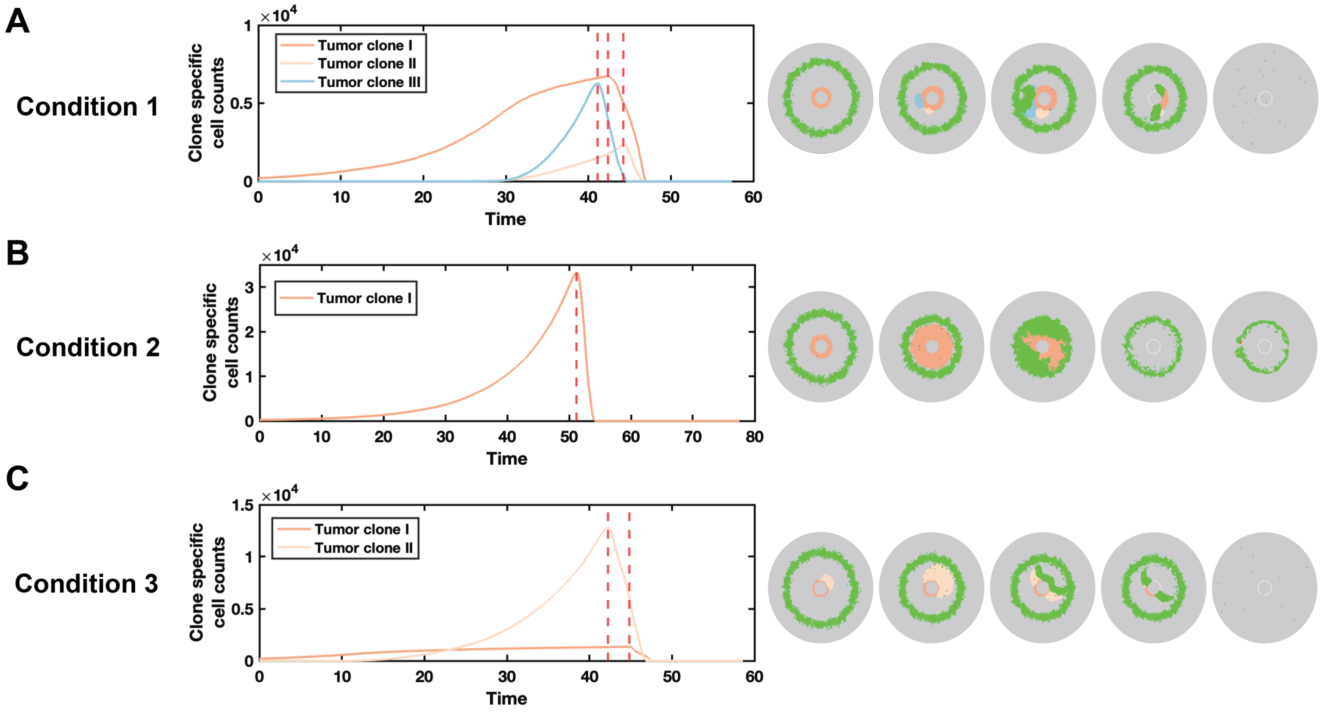


**Fig. S11. The sequence of tumor clone elimination depends on the relative fitness of adjacent clones.** Cell counts for specific clones are shown for conditions 1, 2, and 3. In condition 1, the blue clone demonstrates the highest growth rate. Consequently, the blue clone is the first to be eliminated, followed by the larger orange clone and then the pink clone. Although there are slight differences in growth rates among these clones during this occasion, conditions 2 and 3 reveal instances where a particular clone exhibits a significantly higher growth rate, surpassing all others. These clones become vulnerable to elimination while protecting other nested clones, delaying their immediate elimination. Clones temporarily shielded by others face two potential outcomes: if recognized by T cells, they will be eliminated; conversely, if they find unoccupied space nearby, they can persist in growth and evolution, ultimately evading surveillance.

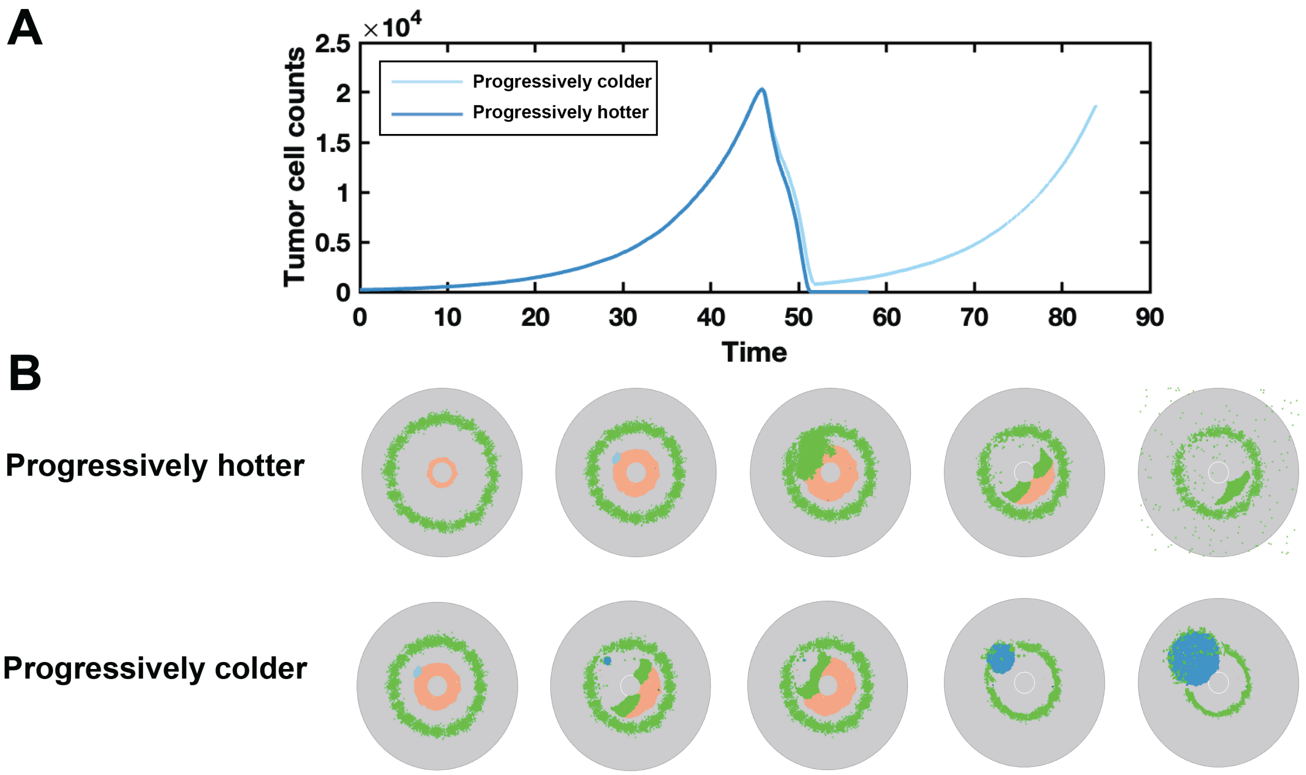


**Fig. S12. The progressively colder tumor presents an escalating challenge for immune killing.** We simulated two conditions and representative examples are shown above: one with increasing immunogenicity, termed "hot", and the other with decreasing immunogenicity, termed "cold". Tumors exhibiting decreasing immunogenicity pose greater difficulties for immune cell infiltration, rendering them more prone to relapse.
